# Toxic Small Alarmone Synthetase FaRel2 inhibits translation by pyrophosphorylating tRNA^Gly^ and tRNA^Thr^

**DOI:** 10.1101/2024.07.05.602228

**Authors:** Tatsuaki Kurata, Masaki Takegawa, Takayuki Ohira, Egor A. Syroegin, Gemma C. Atkinson, Marcus J.O. Johansson, Yury S. Polikanov, Abel Garcia-Pino, Tsutomu Suzuki, Vasili Hauryliuk

## Abstract

Translation-targeting toxic Small Alarmone Synthetases (toxSAS) are effectors of bacterial Toxin-Antitoxin systems that pyrophosphorylate the 3’-CCA end of tRNA to prevent aminoacylation. toxSAS are implicated in antiphage immunity: phage detection triggers the toxSAS activity to shut down viral production. We show that the toxSAS FaRel2 inspects the tRNA acceptor stem to specifically select tRNA^Gly^ and tRNA^Thr^. The 1^st^, 2^nd^, 4^th^ and 5^th^ base pairs the stem act as the specificity determinants. We show that the toxSASs PhRel2 and CapRel^SJ46^ differ in tRNA specificity from FaRel2, and rationalise this through structural modelling: while the universal 3’-CCA end slots into a highly conserved CCA recognition groove, the acceptor stem recognition region is variable across toxSAS diversity. As phages use tRNA isoacceptors to overcome tRNA-targeting defences, we hypothesise that highly evolvable modular tRNA recognition allows for the escape of viral countermeasures through tRNA substrate specificity switching.

## INTRODUCTION

Toxin-antitoxin (TA) systems are ubiquitous prokaryotic regulatory systems. When active, the toxin abolishes bacterial growth – and its toxicity can be efficiently countered by the antitoxin, which can be either RNA-or protein-based^1^. The most common type of TA is Type II, where the protein toxin is neutralised by a protein antitoxin through formation of a tight complex. While multiple biological functions have been attributed to TAs, in recent years it has become clear that many are antiphage defence systems that act through abortive infection mechanisms^2^. TA effectors employ numerous mechanisms of toxicity, often compromising translation^3^. As an essential component of the translational machinery, tRNAs are targeted by many different TA toxin families^4^. To compromise translation, tRNAs can be cleaved by PIN-containing toxins such as VapC^5–8^ and MazF^9^ or enzymatically modified by nucleotidyltransferases such as MenT^10^, toxic Small Alarmone Synthetase (toxSAS) pyrophosphokinases such as FaRel2, CapRel, PhRel and PhRel2^11–13^ or GCN5-related N-acetyl-transferase (GNAT) acetyltransferases such as TacT^14^, AtaT^15–17^ and ItaT^18^.

The substrate specificity of tRNA-targeting TA toxins has been an active topic of research. tRNA-cleaving PIN toxins are typically highly specific, with different members of the same family targeting different tRNA species^5–9^. Substrate specificity of GNAT acetyltransferase toxins that catalyse aminoacyl-tRNA acylation varies from narrow (as in the case of AtaT2 from *Escherichia coli* O157:H7 that exclusively targets Gly-tRNA^Gly^, ref.^17^; or *E. coli* ItaT that specifically targets Ile-tRNA^Ile^, ref.^18^) to relatively broad (*E. coli* AtaT modifies Gly-tRNA^Gly^, Trp-tRNA^Trp^, Tyr-tRNA^Tyr^, Phe-tRNA^Phe^ and Met-tRNA_i_^fMet^, ref.^16^; and a similarly broad specificity was reported for *Salmonella Enteritidis* and *Salmonella Typhimurium* TacTs^19^). The nucleotidyltransferase MenT is particularly specialised. This toxin preferentially adds pyrimidines to the 3’-CCA end of just one tRNA species, tRNA^Ser^, ref.^10^ Phages exploit the tRNA specificity of tRNA-targeting defences by expressing tRNA isoacceptors that are not recognised as substrates, thus allowing translation to continue. This strategy has been observed in the case of T5 phages that resist Retron^20^ and PARIS^21^ defence systems.

toxSASs are effectors of a recently discovered group of Type II TA systems. These toxins are members of RelA/SpoT Homolog (RSH) superfamily, and carry a catalytic domain – toxSYNTH – related to the (pp)pGpp alarmone synthetase domain^22,23^. toxSASs employ two distinct mechanisms of growth inhibition. *Cellulomonas marina* FaRel and the *Pseudomonas aeruginosa* Type VI effector Tas1 synthesise the toxic alarmone (pp)pApp, which, in turn, causes depletion of ATP^12,24,25^. FaRel2, CapRel, PhRel and PhRel2 toxSAS subfamilies on the other hand act as specific inhibitors of protein synthesis, pyrophosphorylating the 3’-CCA end of deacylated tRNA with ATP serving as a pyrophosphate donor^11–13^. FaRel2 from *Сoprobacillus* sp. D7 (recently renamed in NCBI databases to *Thomasclavelia ramose*) forms a non-toxic, enzymatically inert hetero-tetrameric complex with its cognate antitoxin ATfaRel2^26^. Prophage-encoded translation-targeting toxSAS TAs provide narrow-spectrum defence against superinfecting lytic phages, as it was shown for fused TA CapRel^SJ46^, ref^13^, and bipartite TA PhRel:ATphRel (gp29:gp30)^27^. While *Сoprobacillus* sp. D7 FaRel2:ATfaRel2 fails to afford antiphage protection when expressed in *E. coli* and tested against the BASEL phage collection^28^ (**Supplementary Fig. 1a**), this is likely due to the use of a highly heterologous experimental system.

In our initial report describing tRNA pyrophosphorylation by toxSASs, we observed that FaRel2 is tRNA substrate-specific: in biochemical assays the enzyme modifies *E. coli* tRNA^Phe^ more efficiently than tRNA_i_^fMet^, and is exceedingly inefficient in pyrophosphorylating tRNA^Val^, ref.^11^ All of the three tested tRNAs belong to type I as per Brennan-Sundaralingam classification^29^. Type I tRNAs have a short variable loop, while type II tRNAs tRNA^Ser^, tRNA^Tyr^ and tRNA^Leu^ have a long (>10 nt) variable loop with a helical stem of 3-7 base pairs. Structural modeling suggests that the acceptor stem contains the primary substrate specificity determinants recognised by FaRel2^11^. Our original docking model of FaRel2 complexed with *E. coli* tRNA^Phe^ suggested that the 3’-CCA end of tRNA is guided into the FaRel2 active site through multiple contacts with the tRNA acceptor stem^11^. The acceptor stem recognition is mediated by a basic patch of FaRel2, with K28A and R29A substitutions significantly reducing toxicity^11^. In the neutralised FaRel2_2_:ATfaRel2_2_ TA complex, tRNA substrate binding by FaRel2 is seemingly unaffected; instead, the antitoxin exerts its neutralising activity by precluding the binding of the ATP substrate^26^. However, the binding experiments were performed with only one tRNA species, *E. coli* tRNA_i_^fMet^, yielding a K*_D_* of 0.5 μM for both free FaRel2 and its antitoxin-naturalised form^26^. The full *in vivo* substrate specificity of FaRel2 is unknown, and it is similarly unclear whether the tRNA binding specificity of FaRel2 is affected by TA complex formation.

In this study we establish the tRNA specificity of FaRel2. We demonstrate that the toxin preferentially binds and modifies type I tRNAs tRNA^Gly^ and tRNA^Thr^. Both the free FaRel2 and the neutralised TA complex display the same tRNA specificity. Through structural modelling using AlphaFold 3, ref.^30^, RNA substrate mutagenesis and biochemical assays we establish that four nucleotide base pairs in the acceptor stem of the tRNA determine tRNA substrate selection by FaRel2. We show that two translation-targeting toxSASs, *Bacillus subtilis* la1a PhRel2^11^ and fused toxin-antitoxin CapRel^SJ46^, ref.^13^, differ in tRNA specificity from FaRel2. Structural predictions of toxSAS:tRNA complexes by AF3 combined with conservation analyses provide the structural rational for divergent tRNA specificity amongst toxSAS: while the universal 3’-CCA element is recognized by a conserved positively charged groove, the acceptor stem is recognised by a highly divergent site.

## RESULTS

### Both the FaRel2 toxin and the neutralised FaRel2:ATfaRel2 complex preferentially bind type I tRNAs

To establish the tRNA specificity of FaRel2, we isolated toxin-bound tRNAs through FaRel2 immunoprecipitation and identified the enriched RNA species through modification-induced misincorporation tRNA sequencing (mim-tRNAseq)^31^. For selective isolation of FaRel2 from BW25113 *E. coli* cells we used a C-terminally FLAG_3_-tagged FaRel2 variant since, as we have shown previously, this engineered variant retains wild-type level of toxicity while being efficiently neutralised by the ATfaRel2 antitoxin^25^. To improve the yield of FaRel2-FLAG_3_, we co-expressed it with the Small Alarmone Hydrolase (SAH) PaSpo^SSU5^ from *Salmonella* phage SSU5. We previously showed that this SAH efficiently counteracts the toxic effects of FaRel2 by removing the tRNA modification installed by the toxin^11,25^. Finally, as a specificity control, we constructed a FaRel2 variant defective in tRNA binding. As we showed earlier, individual substitutions K28A and R29A that we predicted to disrupt the recognition of the acceptor stem of the tRNA significantly decrease the toxicity of FaRel2^11^. Here we used a double-substituted K28A R29A FaRel2 protein variant that is non-toxic (**Fig. 1a**).

**Figure 1.**
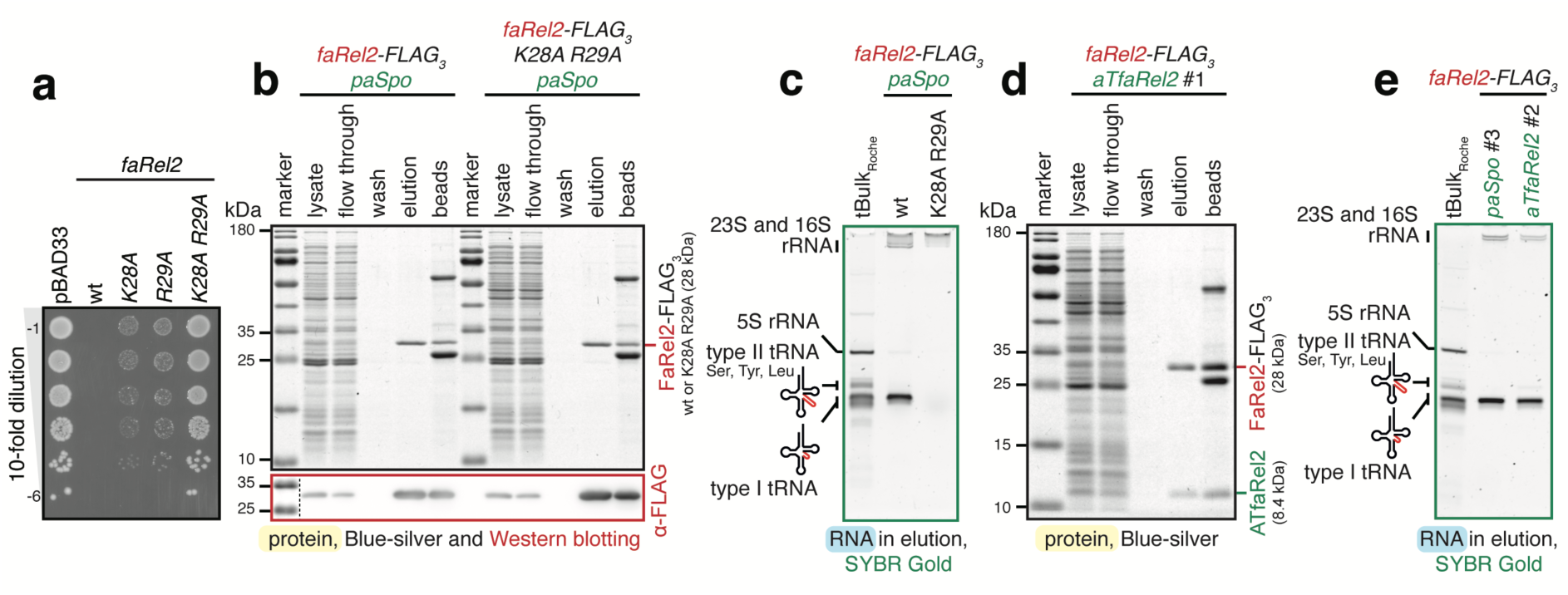
Immunoprecipitated FaRel2 associates with type I tRNAs. **(a)** Ten-fold dilutions of overnight cultures of *E. coli* strains transformed with pBAD33 vector or pBAD33 derivatives expressing either wild-type *Coprobacillus* sp. D7 FaRel2 or alanine-substituted FaRel2 variants: K28A, R29A and the non-toxic double-substituted K28A R29A. **(b)** Purification of C-terminally FLAG_3_-tagged FaRel2 using anti-FLAG-conjugated beads. Samples were separated on SDS-PAGE and visualised by Blue-silver staining as well as by Western blotting with anti-FLAG antibodies. To counter the toxicity of wild-type FaRel2, the toxSAS was co-expressed PaSpo^SSU5^ Small Alarmone Hydrolase (SAH) from *Salmonella* phage SSU5. **(c)** RNA co-eluted with FaRel2-FLAG3 resolved on urea-PAGE and visualised by SYBR Gold staining. **(d)** Immunoprecipitation of FaRel2-FLAG_3_:ATfaRel2 for purification of co-IPed tRNA. **(e)** Comparison of the RNA samples co-eluted with either FaRel2-FLAG_3_ or FaRel2-FLAG_3_:ATfaRel2 with a commercial preparation of *E. coli* small RNA fraction, tBulk_Roche_. Additional replicates are shown on **Supplementary Figure 2**. All experiments were performed at least two times; representative images are shown.

Immunoprecipitated FaRel2-FLAG_3_ and its K28A R29A derivative are highly homogeneous on the protein level (**Fig. 1b**). The RNA component of the FaRel2-FLAG_3_ sample is dominated by tRNA, with minor contamination by rRNA; as expected, no tRNA band is detectable in the case of the K28A R29A variant (**Fig. 1c**). Sucrose gradient centrifugation and immunoblotting shows that FaRel2 does not stably associate with 70S ribosomal complexes, further supporting that the co-IPed tRNAs are directly bound to the toxin (**Supplementary Fig. 1b**). Comparison with the total small RNA sample from *E. coli* revealed that FaRel2-FLAG_3_ specifically coprecipitates with type I – but not type II – tRNA species. Note that in the following experiments we used both a commercial product from Roche, tBulk_Roche_ (used as electrophoresis marker and in enzymatic assays), as well as our own tRNA preparations, designated simply as tBulk (used in enzymatic assays and as a control for mim-tRNAseq, see below).

After establishing the specificity of our pulldown procedure, we immunoprecipitated two types of samples for mim-tRNAseq: i) FaRel2-FLAG_3_ co-expressed with PaSpo^SSU5^ SAH and ii) FaRel2-FLAG_3_ co-expressed with ATfaRel2 antitoxin. The latter preparation contained sub-stochiometric amounts of the antitoxin (**Fig. 1d**). Just as FaRel2-FLAG_3_, FaRel2-FLAG_3_:ATfaRel2 coprecipitated with type I tRNAs (**Fig. 1e**). Finally, we re-purified the tRNA fractions from rRNA contamination through sizing on a urea-PAGE gel and used the resultant samples for mim-tRNAseq library preparation (≥1µg tRNA per sample; replicates and controls are shown on **Supplementary Fig. 2**).

### FaRel2 and FaRel2:ATfaRel2 preferentially bind tRNA^Gly^ and tRNA^Thr^

After deacylation of tBulk and immunoprecipitated tRNA samples, DNA adapters were ligated to tRNA CCA 3’ ends, the ligated RNA-DNA products gel-purified, subjected to reverse transcription and resolved on urea-PAGE. The cDNA ran as two bands, one being a full-length product and the other being an aberrant product caused by RT-stop due to tRNA modifications (**Supplementary Fig. 3**). Both cDNA species were gel-purified, circularized and PCR-amplified. The resultant cDNA library was gel-purified once again before Illumina sequencing. The reads were processed and aligned to non-coding *E. coli* RNA genomic sequences. Taking advantage of 17 nucleotide random UMI (D3N14) barcodes at the 5’ end of the RT primers, duplicated reads were removed after genome mapping, and the remaining reads were counted for individual tRNA genes (**Supplementary Table 1**).

tRNA pools that co-purify with both FaRel2 and FaRel2:ATfaRel2 are dominated by type I tRNAs tRNA^Gly^ and tRNA^Thr^ (**Figure 2a-c**). In the case of tRNA^Gly^, the FaRel2-and FaRel2:ATfaRel2-coimmunoprecipitated samples were strongly enriched in tRNA^Gly1^ (6-and 7-fold enrichment, respectively), tRNA^Gly2^ (11-and 12-fold) and tRNA^Gly3^ (6-and 7-fold). Strong enrichment was observed for tRNA^Thr1^ and tRNA^Thr3^ (both 7-fold), as well as a modest enrichment for tRNA^Thr4^ (2-fold). The fraction of the tRNA^Thr2^ remains largely unchanged, which is indicative of a higher affinity than the majority of tRNA species which were depleted during immunoprecipitation. Despite being more abundant in tBulk, tRNA^Thr4^ (2-fold enrichment) is less dominant in the immunoprecipitated pool as compared to tRNA^Gly2^ (11-and 12-fold enrichment), suggesting more efficient ‘capture’ of tRNA^Gly2^ by FaRel2 in the cell. Notably, the three tRNA species that were used for biochemical studies of FaRel2 earlier – tRNA^Phe^, tRNA_i_^fMet^ and tRNA^Val^, refs.^11,26^ – were virtually absent in the immunoprecipitated tRNA pool. This, however, does not mean that FaRel2 has no affinity to these tRNA species (it was earlier established that FaRel2 binds *E. coli* tRNA_i_^fMet^ with a K_D_ of 0.5 μM, ref.^26^); rather, the weaker binders are outcompeted by the tighter binders.

**Figure 2.**
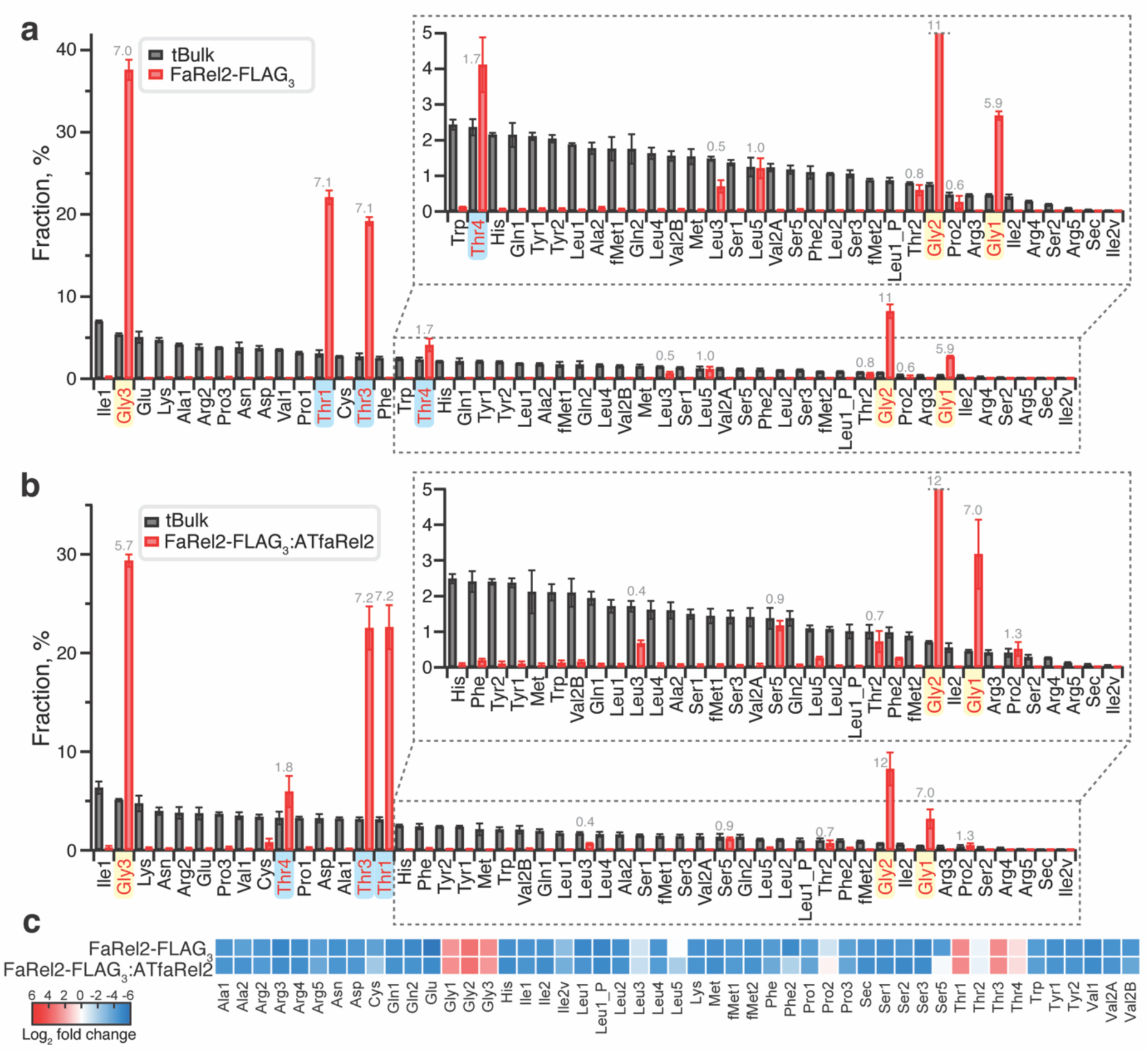
tRNA^Gly^ and tRNA^Thr^ are specifically bound by FaRel2 and FaRel2:ATfaRel2. **(a,b)** tRNA species abundance in **(a)** FaRel2-FLAG_3_-or **(b)** FaRel2-FLAG_3_:ATfaRel2-coimmunoprecipitated RNA fraction as well as *E. coli* tBulk was quantified by mim-tRNAseq^31^. Fold-change of the relative tRNA abundance (fraction in co-immunoprecipitated pool vs tBulk) is shown as grey numbers. **(c)** Heat map representation for log_2_-fold change in abundance of tRNA species in coimmunoprecipitated tRNA samples relative to their abundances in tBulk. The experiments were performed at least three times; the data are shown as average ± standard deviation.

### FaRel2 abrogates translation by preferentially targeting tRNA^Gly^ and tRNA^Thr^

While the immunoprecipitation-based assays are informative in establishing the binding specificity of FaRel2, they do not necessarily reflect the enzymatic substrate preferences of the toxin. To address this question, we first assessed the effects of FaRel2 expression on tRNA_i_^fMet^ and tRNA^Thr1^ charging levels via Northern blotting assays. Consistent with the inferred substrate specificity, expression of FaRel2 resulted in the disappearance of the acylated tRNA^Thr1^ without affecting the charging levels of tRNA_i_^fMet^ (**Figure 3a**).

**Figure 3.**
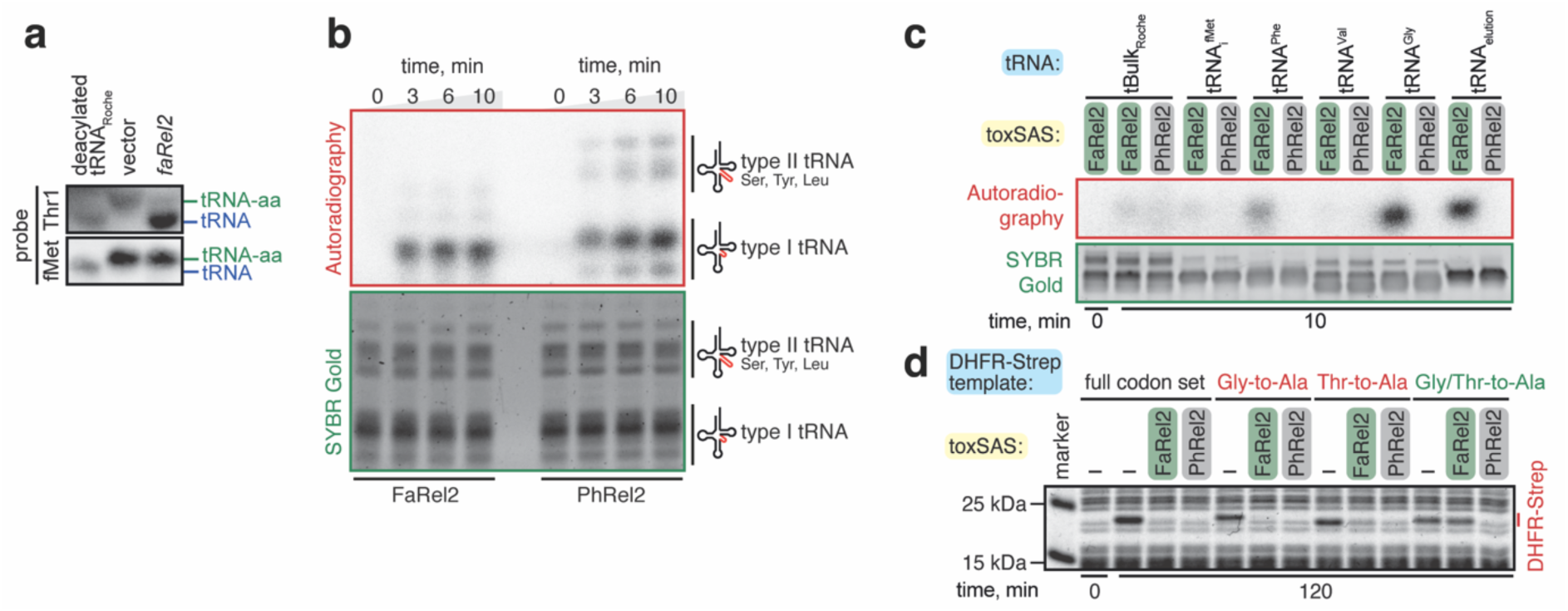
FaRel2 inhibits translation by specifically pyrophosphorylating tRNA^Gly^ and tRNA^Thr^. **(a)** Total RNA prepared from cells either expressing FaRel2 or transformed with an empty vector plasmid was resolved on acidic urea-PAGE and probed against tRNA^Thr1^ or tRNA_i_^fMet^. **(b,c)** tRNA pyrophosphorylation by FaRel2 or *B. subtilis* la1a PhRel2 assayed using ^32^P-labelled ATP. **(b)** FaRel2 specifically labels the type I tRNAs from *E. coli* total tRNA, tBulk_Roche_. 50 nM toxSAS-FLAG_3_ was reacted with 5 μM tBulk_Roche_ at 37°C for increasing periods of time. tBulk_Roche_ stands for commercial preparation of *E. coli* small RNA fraction. **(c)** RNA^Gly^ and FaRel2-FLAG3-coimmunoprecipitated tRNA fractions are more efficiently modified by FaRel2 as compared to tBulk_Roche_ and individual *E. coli* tRNAs tRNA_i_^fMet^, tRNA^Phe^ and tRNA^Val^. 50 nM toxSAS-FLAG_3_ was reacted with 0.4 μM of tRNA preparations at 37°C for 10 min. **(d)** Reporter expression assays in cell-free protein synthesis system. Addition of FaRel2 abrogates production of Strep-tagged DHFR reporter proteins which harbours full codon set or in which all Gly or Thr codons were substituted to Ala (Gly-to-Ala or Thr-to-Ala), but not the mutant variant in which all Gly and Thr codons were converted to Ala (Gly/Thr-to-Ala). Addition of *B. subtilis* la1a PhRel2 abrogates the expression of all reporters equally. All experiments were performed at least two times; representative gels are shown.

Next, we performed an *in vitro* tRNA pyrophosphorylation assay with FaRel2 using *E. coli* tBulk_Roche_ and ^32^P-ATP as substrates^11^. In good agreement with the immunoprecipitation experiments, the majority of the ^32^P signal comes from type I tRNA species (**Figure 3b**). As a specificity control, we performed the labelling assay with another translation-targeting toxSAS, *B. subtilis* la1a PhRel2^11^. In the case of PhRel2, the labelling pattern is different (**Figure 3b**). Both type I and type II tRNAs are modified, and in the case of type I tRNA, both low and high-Mw species are labelled. Importantly, when tested under same conditions, FaRel2 and PhRel2 were similarly efficient.

To probe the enzymatic specificity of FaRel2 further, we then performed pyrophosphorylation assays with either i) tBulk, ii) native *E. coli* tRNA_i_^fMet^, tRNA^Phe^, tRNA^Val^ and tRNA^Gly^ and, finally iii) tRNA^Gly^-and tRNA^Thr^-enriched tRNA fractions isolated through anti-FLAG_3_ immunoprecipitation of FaRel2-FLAG_3_. Just as in the previous experiment, we used PhRel2 for comparison, and, in order to increase the selectivity of tRNA modification, the tRNA substrate was used at lower concentration as compared to the previous experiment (0.4 µM vs 5 µM). tRNA co-IPed with either FaRel2-FLAG_3_ or FaRel2-FLAG_3_:ATfaRel2 was modified by FaRel2 more efficiently than either tBulk or tRNA_i_^fMet^; tRNA^Gly^ was modified as efficiently as the co-IPed tRNA (**Figure 3c** and **Supplementary Figure 4**). In the case of PhRel2, none of the tested tRNA species and fractions were modified efficiently (**Figure 3c**), suggesting tRNA_i_^fMet^, tRNA^Phe^, tRNA^Val^, tRNA^Gly^ and tRNA^Thr^ are not modified by PhRel2 efficiently.

Taken together, our results establish that i) the preferential binding of tRNA^Gly^ and tRNA^Thr^ by FaRel2 is, indeed, reflective of the toxin’s enzymatic preferences and ii) the substrate preferences are toxSAS-specific, as PhRel2 clearly has a different specificity. We reasoned that, given FaRel2’s strong preference of these two specific tRNA species, expression of proteins that do not contain glycine or threonine should be largely insensitive to FaRel2. To test this, we used the reconstituted cell-free protein synthesis system from *E. coli* components (PURE)^32^. As we showed earlier, FaRel2 efficiently inhibits production of dihydrofolate reductase (DHFR) in the PURE system^11^. We constructed four variants of the Strep-tagged DHFR reporter. The first variant was designed to contain the full set of 61 sense codons (designated as full codon set). The second and third variants were based on the full set reporter, but all of either glycine or threonine-encoding codons are substituted for alanine (designated as Gly-to-Ala and Thr-to-Ala, respectively). Finally, we constructed a variant in which both glycine and threonine-encoding codons are substituted for alanine, GT-to-A. All reporters described above were tested with FaRel2, PhRel2 and the fused toxin-antitoxin CapRel^SJ46^, ref^13^. The latter enzyme is inactive unless triggered by the SECΦ27 phage major capsid protein Gp57^13^. Directly supporting our predictions, while the expression of the full codon set DHFR reporter as well as its Gly-to-Ala and Thr-to-Ala variants is readily abrogated by FaRel2, expression of the Gly/Thr-to-Ala variant is insensitive to FaRel2 (**Figure 3e**). Both PhRel2 (**Figure 3e**) and Gp57-triggered CapRel^SJ46^ (**Supplementary Figure 5**) efficiently inhibit the synthesis of all reporters, suggesting that these toxSAS toxins display a tRNA substrate specificity that is different from that of FaRel2.

### toxSASs combine a conserved CCA-recognising groove with divergent substrate specificity regions

To gain a structural insight into toxSAS tRNA selectivity, we predicted the structures of tRNA^Gly1^-bound FaRel2, PhRel2 and CapRel^SJ46^ in the ATP-liganded state using AlphaFold 3 (AF3), ref.^30^. We plotted charge distribution (**Figure 4a-c**) and conservation as calculated by ConSurf^33^ (**Figure 4d-f**) on the protein structures. As we do not know which tRNA species are preferred by PhRel2 and CapRel^SJ46^, in order to simplify comparison, the same tRNA^Gly1^ was used for all predictions. The AF3-generated model of FaRel2:tRNA^Gly1^ is consistent with both the crystal structure of FaRel2, ref.^26^, as well as the FaRel2:tRNA docking model generated using HADDOCK^34^ and validated though mutagenesis^11^. Mutational analysis supports the AF3-generated models of tRNA^Gly1^-bound PhRel2 and CapRel^SJ46^ (**Supplementary Figure 6**).

**Figure 4.**
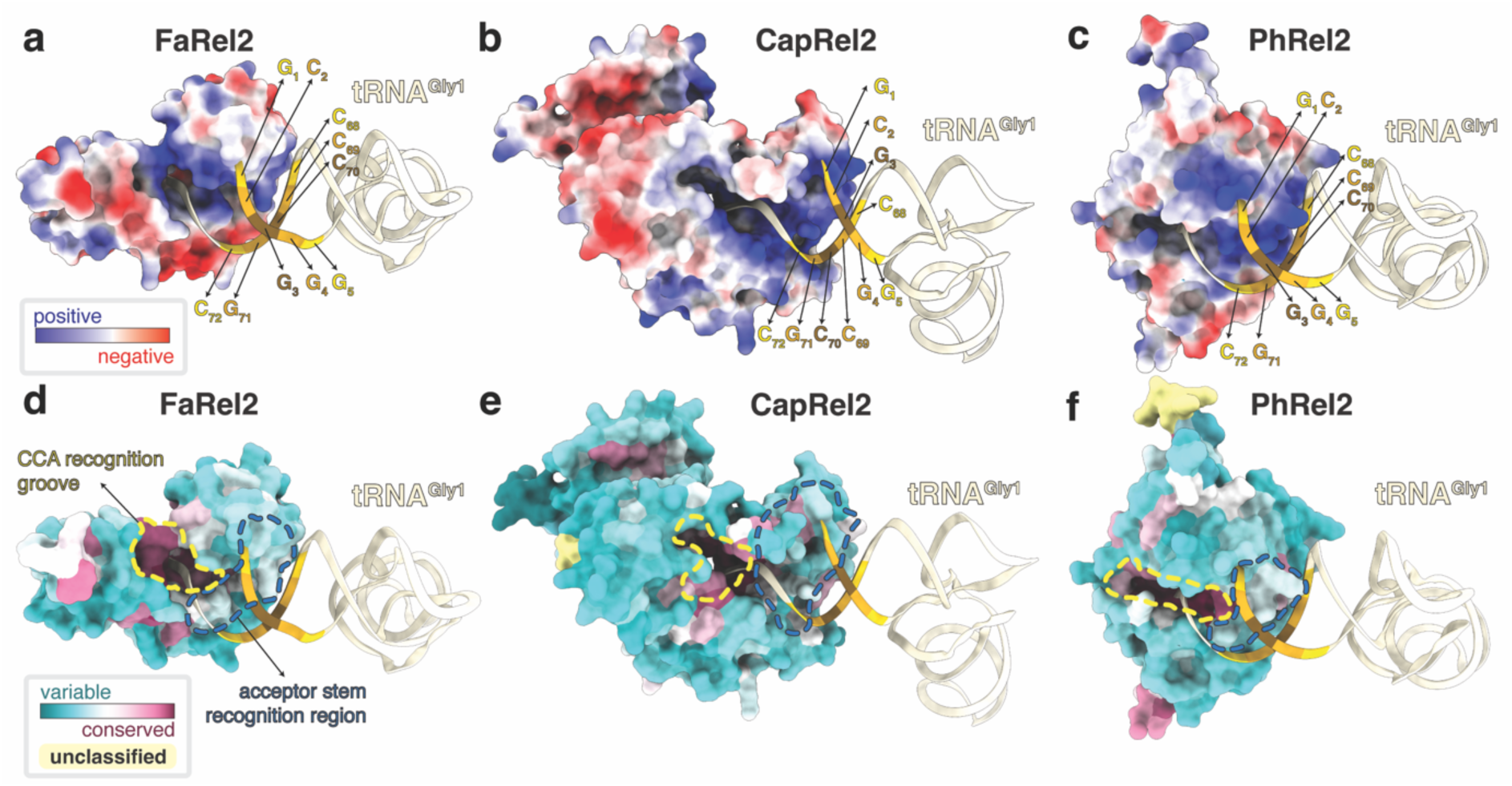
Conservation and diversification of tRNA recognition by toxSAS. **(a-c)** AF3-generated models of tRNA^Gly1^-bound *Coprobacillus* sp. D7 FaRel2, *B. subtilis* la1a PhRel2 and CapRel^SJ46^ coloured by the surface charge. **(d-f)** Same toxSAS:tRNA^Gly1^ complexes as on (a-c), but coloured by amino acid conservation as computed by ConSurf^33^.

The predicted structures universally place the tRNA acceptor stem and the 3’-CCA to be recognised by a positively charged surface of the toxSAS (**Figure 4a-c**). The 3’-CCA is slotted into a deep and highly basic CCA-recognition groove that extends to the toxSYNTH active site, while the first five base pairs of the acceptor stem interact with a shallower but similarly basic acceptor stem recognition region. While the CCA-recognition groove is conserved among toxSASs, the acceptor stem recognition region is divergent, thus explaining the different tRNA substrate specificity amongst FaRel2, PhRel2 and CapRel^SJ46^ (**Figure 4d-f**). *G_1_-C_72_, C_2_-G_71_, G_4_-C_69_ and G_5_-C_68_ serve as positive determinants for selection of tRNA^Gly^ and tRNA^Thr^ by FaRel2* Structural modelling suggests that FaRel2 contacts the first five base pairs of the acceptor stem of the tRNA (**Figure 4**). This suggests that the substrate specificity should be encoded in this region of the tRNA molecule. Sequence comparison amongst tRNA^Gly1^, tRNA^Gly2^, tRNA^Gly3^, tRNA^Thr1^, tRNA^Thr3^ and tRNA^Thr4^ suggests that G_1_-C_72_, C_2_-G_71_, G_4_-C_69_ and Pu_5_-Py_68_ of the acceptor stem could serve as positive determinants for the recognition by FaRel2 since these nucleotides are conserved among the tRNA species that are preferentially recognised by the toxin (**Figure 5a**). The third base pair is unconserved (**Figure 5a**), and, therefore, is unlikely to be specifically recognised by FaRel2. Importantly, while our analyses of FaRel2 tRNA specificity were performed in an *E. coli* surrogate host, tRNA^Gly^, tRNA^Thr^ and tRNA^Leu^ of the *Coprobacillus* native host do contain all the predicted acceptor stems determinants (**Supplementary Table 2**).

**Figure 5.**
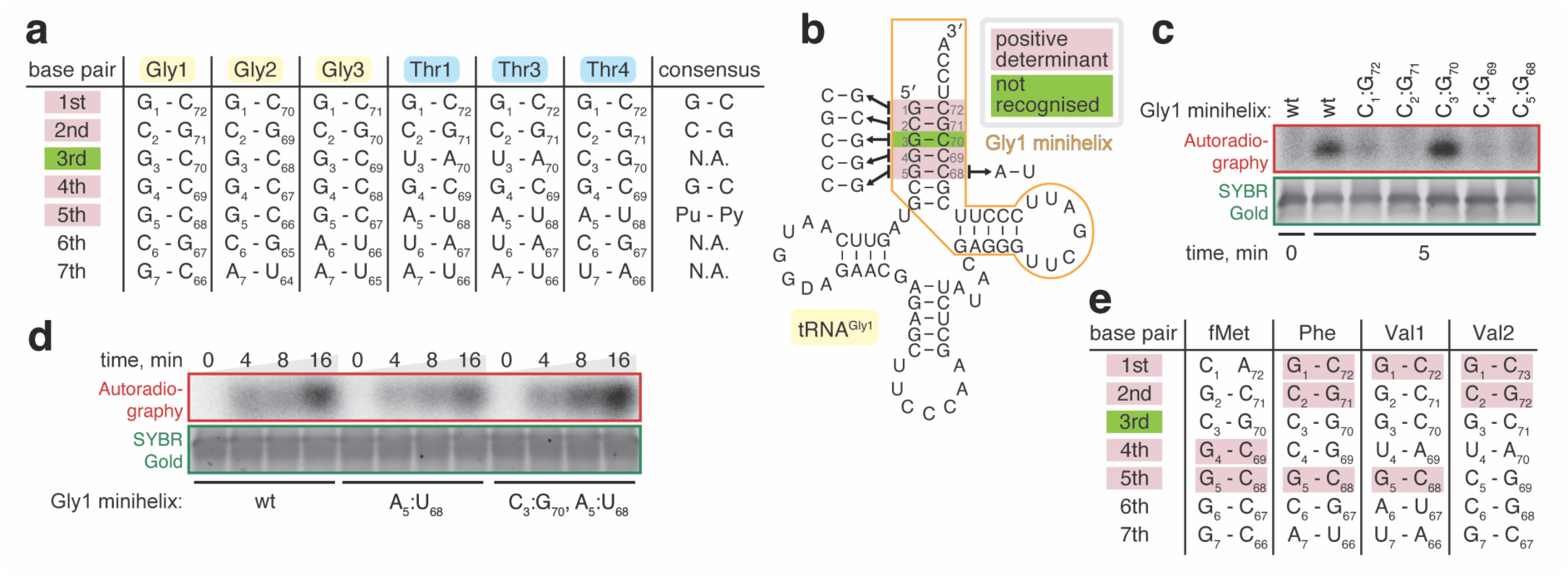
Molecular determinants defining tRNA selection by FaRel2. **(a)** Acceptor stem sequences for *E. coli* tRNA^Gly^ and tRNA^Thr^ isoacceptors. **(b)** *E. coli* tRNA^Gly1^ secondary structure and the tested base pair swapping mutations. The minihelix part is outlined with an orange line. Base pair swapping mutations are indicated by arrowheads with mutation names (1:72, 2:71, 3:70, 4:69 and 5:68). Pink background shows the base pairs where swapping mutation abrogates pyrophosphorylation of mini helix by FaRel2 and green background indicates the base pair where the swapping mutation did not decrease the pyrophosphorylation. **(c)** Pyrophosphorylation of tRNA^Gly1^-mimicking RNA minihelix by FaRel2 assayed with ^32^Plabelled ATP. Base pairs at positions 1-72, 2-71, 4-69 and 5-68 but that at 3-70 are crucial for substrate recognition by FaRel2. 5 nM FaRel2-FLAG_3_ was reacted with 5 μM tRNA minihelix at 37°C for 5 min. The experiments were performed at least three times, representative gels are shown. **(d)** Kinetic analysis of FaRel2-medited modification of wild-type as well as A_5_:U_68_ and A_5_:U_68_ C_3_:G_70_ variants of tRNA^Gly1^-mimicking RNA minihelix. The experiments were performed analogously to (c). **(e)** Acceptor stem sequences for *E. coli* tRNA_i_^fMet^, tRNA^Phe^, tRNA^Val1^ and tRNA^Val2^.

To test the functional importance of these candidate positive specificity determinants, we used synthetic minihelix RNA oligonucleotides mimicking tRNA^Gly1^, a wild-type version as well as a series of base-flipping variants targeting the determinants such as G_1_-C_72_ to C_1_-G_72_ (**Figure 5b**). Using native *E. coli* tRNA^Gly^ as a positive control, we have validated that our tRNA^Gly1^-mimicking RNA minihelix is efficiently modified by FaRel2, thus establishing the validity of our experimental system (**Supplementary Figure 7a**). Base-flipping of either of the candidate determinants – but not of the C_3_-G_70_ pair that is not conserved amongst the substrate tRNA species – compromised minihelix modification by FaRel2 (**Figure 5c**). These results are consistent with the AF3-generated FaRel2:tRNA^Gly1^ model suggesting the lack of strong contacts between the C_3_-G_70_ base pair and the specificity region of FaRel2 (**Figure 4a**). Our kinetically resolved experiments failed to detect any deleterious effect of substitutions of the C_3_-G_70_ pair, further supporting that the base moieties of these nucleotides are not inspected by FaRel2 (**Supplementary Figure 7b,c**). While the substitutions in G_1_-C_72_ decreased the efficiency of modification drastically, this mutant was pyrophosphorylated more efficiently than those targeting C_2_-G_71_, G_4_-C_69_ and G_5_-C_68_, suggesting its relatively lower importance is substrate recognition (**Supplementary Figure 7c**). The A_5_-U_68_ minihelix variant (this base pair is present in all the tRNA^Thr^ isoacceptors) was also efficiently modified, which is consistent with the conservation of this base pair in enriched tRNA species (**Figure 5d**). The A_5_-U_68_ C_3_-G_70_ double substitution that mimics tRNA^Thr2^ and tRNA^Thr4^ did not change the modification efficiency (**Figure 5d**), despite the relatively lower enrichment of these isoacceptors in co-IPed tRNA pools as compared to tRNA^Thr1^ and tRNA^Thr3^. Collectively, our results establish the critical role of G_1_-C_72_, C_2_-G_71_, G_4_-C_69_ and Pu_5_-Py_68_ in tRNA^Gly^ substrate recognition by FaRel2.

## Discussion

The tRNA acceptor stem is recognised by multiple tRNA-binding proteins, such as aminoacyl-tRNA synthetases^35^, ProXp-ala^36^ trans-editing protein that catalyses the hydrolysis of mischarged Ala-tRNA^Pro^, peptidyl-tRNA hydrolase (Pth)^37^, elongation factors EF-Tu/eEF1A^38^ as well diverse toxins such as aminoacyl-tRNA acetylating AtaT^16^, TacT^39^ and ItaT^40^; see recent review by Zhang^41^. Here we dissect one more class of acceptor stem-recognising proteins: translation-targeting toxSAS toxins.

We show that the *Coprobacillus* sp. D7 FaRel2 toxin specifically recognises and pyrophosphorylates tRNA^Gly^ and tRNA^Thr^ and establish that G_1_-C_72_, C_2_-G_71_, G_4_-C_69_ and Pu_5_-Py_68_ base pairs of the tRNA acceptor stem serve as specificity determinants that guide tRNA selection. These insights allow us to rationalise our previous biochemical results^11^, specifically that *E. coli* tRNA^Phe^ is modified more efficiently than initiator tRNA_i_^fMet^ while tRNA^Val^ is an extremely poor substrate: while tRNA^Phe^ contains three determinants, both tRNA_i_^fMet^ and tRNA^Val^ contain only two. Specifically, tRNA^Phe^ has G_1_-C_72_, C_2_-G_71_ and G_5_-C_68_, tRNA_i_^fMet^ has G_4_-C_69_ and G_5_-C_68_, tRNA^Val^ isoacceptors have G_1_-C_72_ as well as either C_2_-G_71_ or G_5_-C_68_ (**Figure 5e**). Finally, we show that *B. subtilis* la1a PhRel2 and CapRel^SJ46^ possess tRNA specificity different from that of FaRel2, and rationalise this observation by structural modelling and conservation analysis.

Our AF3-generated structural models suggest that tRNA selection by toxSAS is mediated by two recognition interfaces. While the universal CCA element is slotted into a highly conserved CCA recognition groove, the tRNA specificity determinants are inspected by the acceptor stem recognition region that is highly variable across the toxSAS diversity (**Figure 4**). As we have shown earlier, even a single-strand 5’-CACCA-3’ RNA pentanucleotide can be modified by FaRel2 if used in a sufficiently high concentration^11^. As this interaction would be mediated exclusively by the CCA recognition groove, it is to be expected that the affinity for this minimalistic substrate is low. The bipartite tRNA recognition mode of toxSAS is similar to that of aminoacyl-tRNA-modifying GNAT toxins such as AtaT^16^ and TacT^39^. Just like the toxSAS, GNAT toxins interact with the CCA via a highly conserved active site patch while “reading” the major groove of the acceptor stem via a variable region. It was shown recently that phages have can overcome tRNA-targeting PARIS^21^ and Retron^20^ immunity systems by supplying alternative phage-encoded tRNA isoacceptors that are not recognised by the defences. Evolutionary plasticity of the acceptor stem recognition region would allow for tRNA specificity switching in diverse toxSAS that would, in turn, enable the escape of these counter-defence measures.

## ACKNOWLEDGMENTS

We are grateful to the Protein Expertise Platform Umeå University and Mikael Lindberg for plasmid construction, and to Alexander Harms for sharing the BASEL phage collection. Computations were partially performed on the NIG supercomputer at ROIS National Institute of Genetics. This work was supported by the Swedish Research Council (Vetenskapsrådet) grants (2019-01085 and 2022-01603 to GCA, 2021-01146 to VH), Crafoord foundation (project grant Nr 20220562 to VH), the Estonian Research Council (PRG335 to VH), Cancerfonden (20 0872 Pj to VH), the project grant from the Knut and Alice Wallenberg Foundation (2020-0037 to GCA and VH), the Fonds National de Recherche Scientifique (FNRS CDR J.0066.21 and J.0065.23F; FNRS-EQP UN.025.19; FNRS PDR T.0066.18 and T.0090.22; and FNRS WELBIO ADV X.1520.24F to AGP); ERC (CoG DiStRes, n° 864311 to AGP); Fonds Jean Brachet and the Fondation Van Buuren (AGP), Exploratory Research for Advanced Technology 1295 from the Japan Science and Technology Agency (JST-ERATO) (JPMJER2002 to TS), the National Institutes of Health (R01-GM132302 to YSP), the National Science Foundation (MCB-1907273 to YSP), and the Illinois State startup funds (YSP).

## Author contributions

TK, GCA, AGP and VH drafted the manuscript with contributions from all authors. TK, TS, AGP and VH coordinated the study. EAS and YSP expressed and purified native *E. coli* tRNA^Gly^. TK, VH, AGP, TS, MJOJ and TO designed experiments, performed experiments and analysed the data. MT and TO performed tRNA-seq and bioinformatic analyses.

## SUPPLEMENTARY TABLES AND FIGURES

**Supplementary Table 1. min-tRNAseq read counts.**

The table is provided as a separate Excel file.

**Supplementary Table 2. Acceptor stem and complete sequences of Coprobacillus sp. D7 tRNAGly, tRNAThr and tRNALeu tRNA species as well as *E. coli* BW25113 complete tRNA sequences.**

The table is provided as a separate Excel file.

**Supplementary Table 3. Strains, plasmids and oligonucleotide primers used in this study.**

The table is provided as a separate Excel file. Tables are in individual tabs with the following information: plasmids, cloning procedures, primers and strains.

**Supplementary Figure 1.**
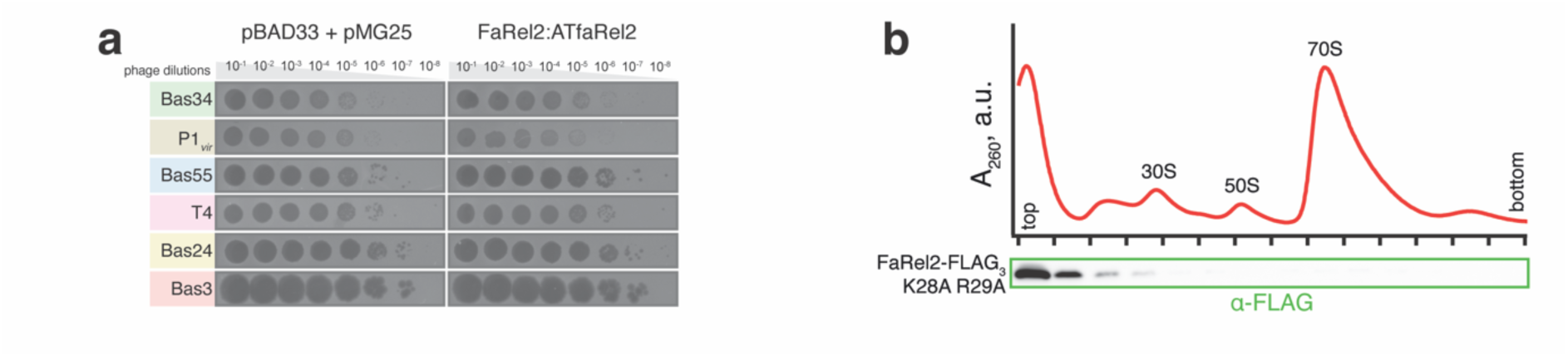
Functional studies of *Coprobacillus* sp. D7 FaRel2:ATfaRel2 TA system. **(a)** *Coprobacillus* sp. D7 FaRel2:ATfaRel2 TA does not afford protection against coliphages when heterologously expressed *E. coli*. Serial dilutions of select BASEL^28^ and common laboratory phages were spotted on a lawn of BW25113 *E. coli*, either expressing FaRel2:ATfaRel2 TA system or transformed with empty pBAD33 and pMG25 plasmids. Experiments with representative phages (out of >60 tested) are shown. **(b)** FaRel2 does not stably associate with ribosomes. A lysate prepared from *E. coli* cells expressing a non-toxic FaRel2 variant (FaRel2-FLAG_3_ K28A R29A) was fractionated on a 10-35% sucrose gradient, and the FLAG_3_-tagged FaRel2 K28A R29A protein was detected by Western blotting with anti-FLAG antibodies.

**Supplementary Figure 2.**
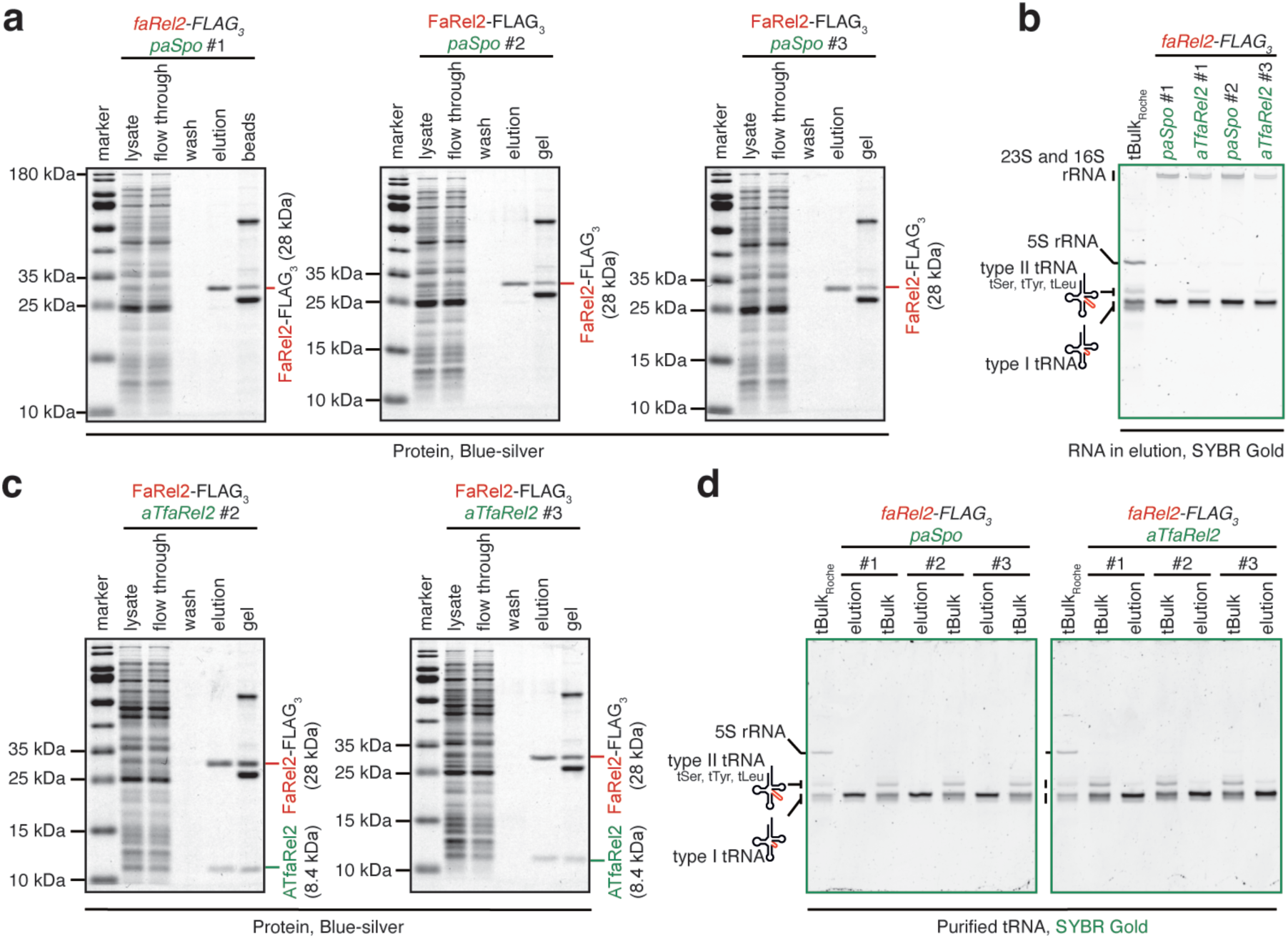
Immunoprecipitation of FaRel2-FLAG_3_ and FaRel2-FLAG_3_:ATfaRel2, two replicates, related to Figure 2. **(a)** Three biological replicates of a-FLAG_3_ immunoprecipitation of FaRel2-FLAG_3_. **(b)** Two additional biological replicates of RNA co-eluted with either FaRel2-FLAG_3_ or FaRel2-FLAG_3_:ATfaRel2 compared to tBulk_Roche_. **(c)** Two replicates of anti-FLAG_3_ immunoprecipitation of FaRel2-FLAG_3_:ATfaRel2. **(d)** Final tRNA fractions used for mim-tRNAseq. tBulk_Roche_ stands for commercial preparation of *E. coli* small RNA fraction, while tBulk designates lab-made preparations.

**Supplementary Figure 3.**
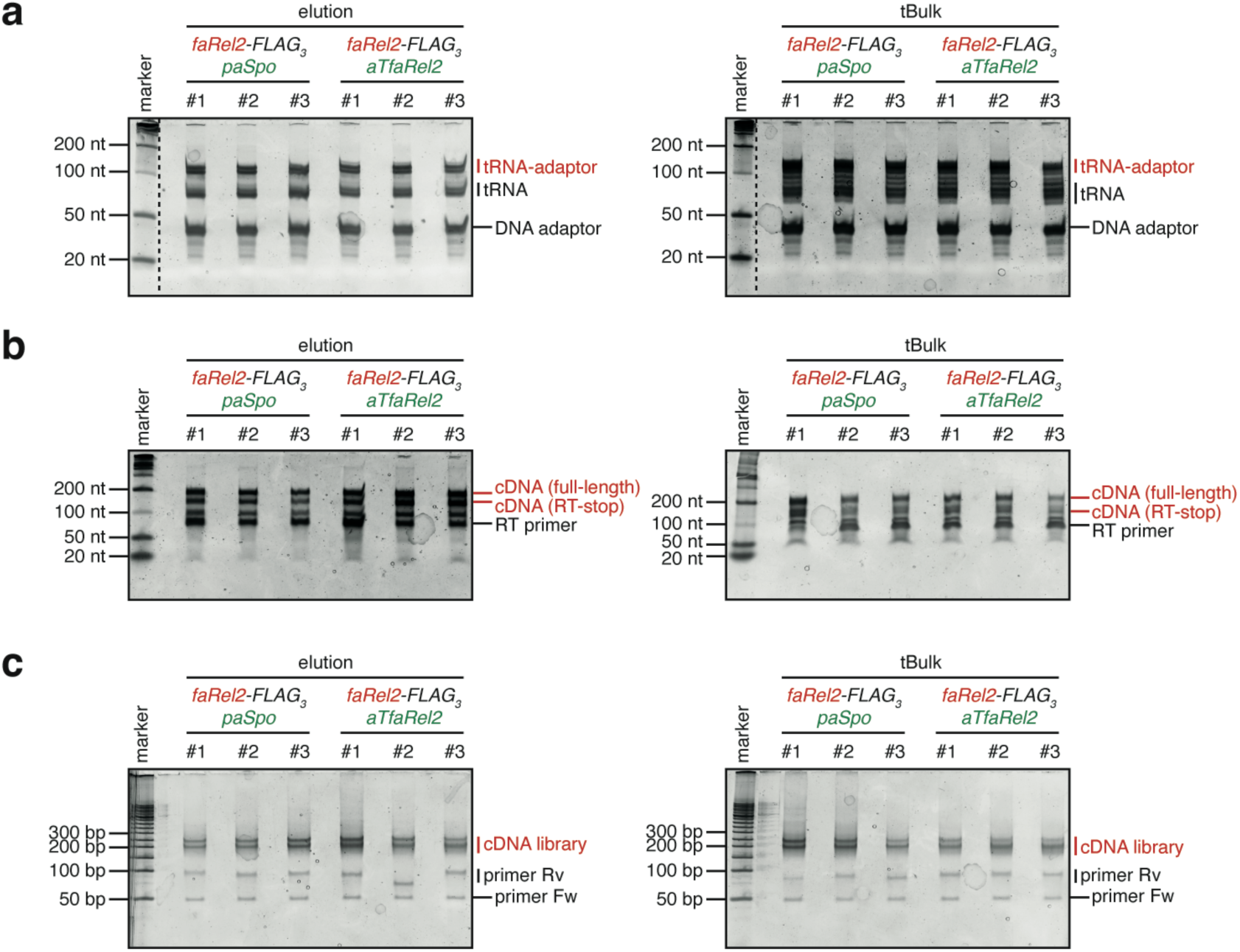
Gel images of samples in the preparation for mim-tRNAseq. Three biological replicates of tBulk preparations and FaRel2-bound tRNA preparations after adaptor ligation **(a)**, reverse transcription **(b)**, and PCR reaction for cDNA library construction **(c)** were resolved on urea-PAGE in 1x TBE (7 M urea, 10% PAGE) or native-PAGE (6% PAGE) in 1x TBE, stained with SYBR Gold, and subjected to gel purification.

**Supplementary Figure 4.**
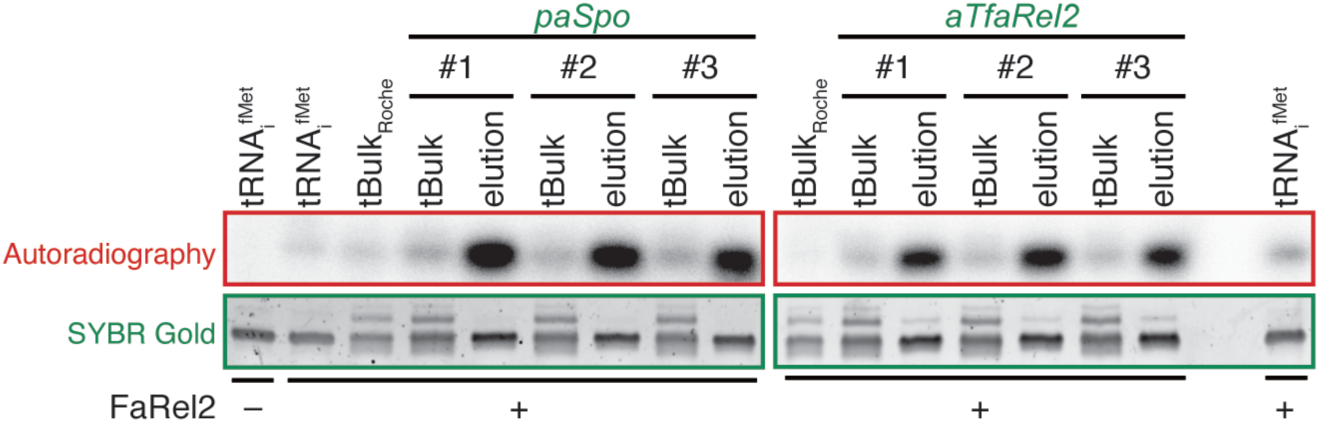
tRNA fraction associated with FaRel2-FLAG_3_:ATfaRel2 is efficiently modified by FaRel2, related to Figure 4. Pyrophosphorylation assays using ^32^P-labelled ATP and unlabelled tRNA substrates. FaRel2-FLAG_3_-or-FLAG_3_:ATfaRel2-coimmunoprecipitated tRNA fractions are more efficiently modified FaRel2 as compared to tBulk and individual *E. coli* tRNA_i_^fMet^. 100 nM FaRel2-FLAG_3_ was reacted with 0.4 µM tRNA substrates at 37°C for 10 min. tBulk_Roche_ stands for commercial preparation of *E. coli* small RNA fraction, while tBulk designates lab-made preparations.

**Supplementary Figure 5.**
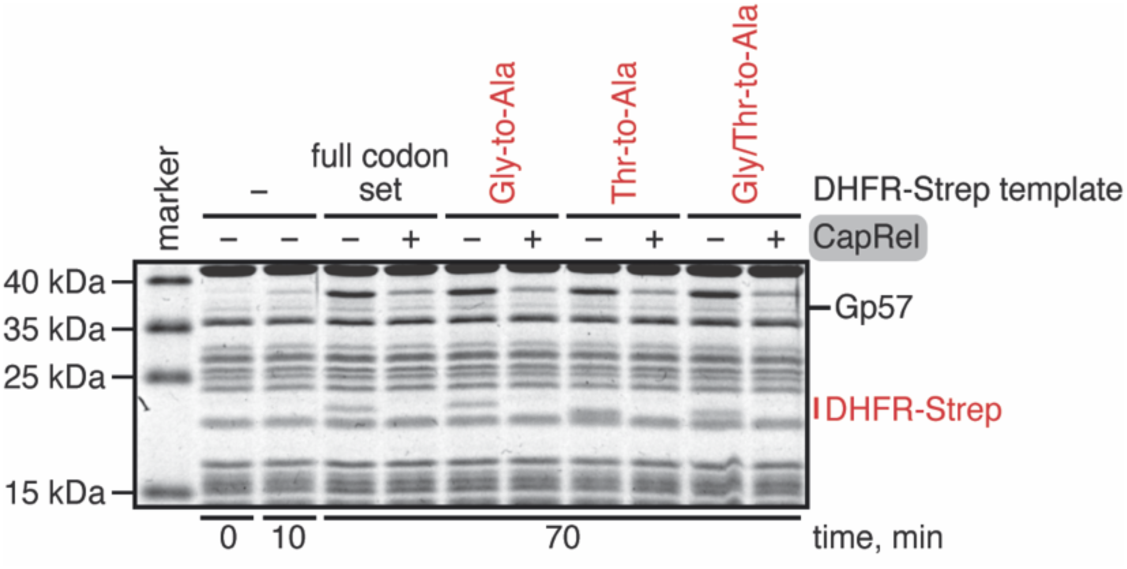
tRNA specificity of CapRel^SJ46^ is different from that of FaRel2. SECΦ27 major capsid protein Gp57, the trigger of CapRel^SJ46^ toxSAS, was produced *in situ* from the template plasmid (10 ng/µl) in the PURE cell-free protein synthesis system. Gp57 was synthesised either in the presence or absence of purified CapRel^SJ46^ (250 nM). Next, the DHFR template plasmids were added (20 ng/µL) and the DHFR reporter proteins were synthesised for 60 minutes at 37°C. The reporters used: i) the full codon set version that encodes all of the possible codons, ii) Gly-to-Ala variant in which all Gly codons substituted for Ala, iii) Thr-to-Ala, all Thr codons substituted for Ala, iv) Gly/Thr-to-Ala, all Gly and Thr codons substituted for Ala. Addition of CapRel^SJ46^ abrogated the production of all of the tested Strep-tagged DHFR reporters. The experiment was performed once.

**Supplementary Figure 6.**
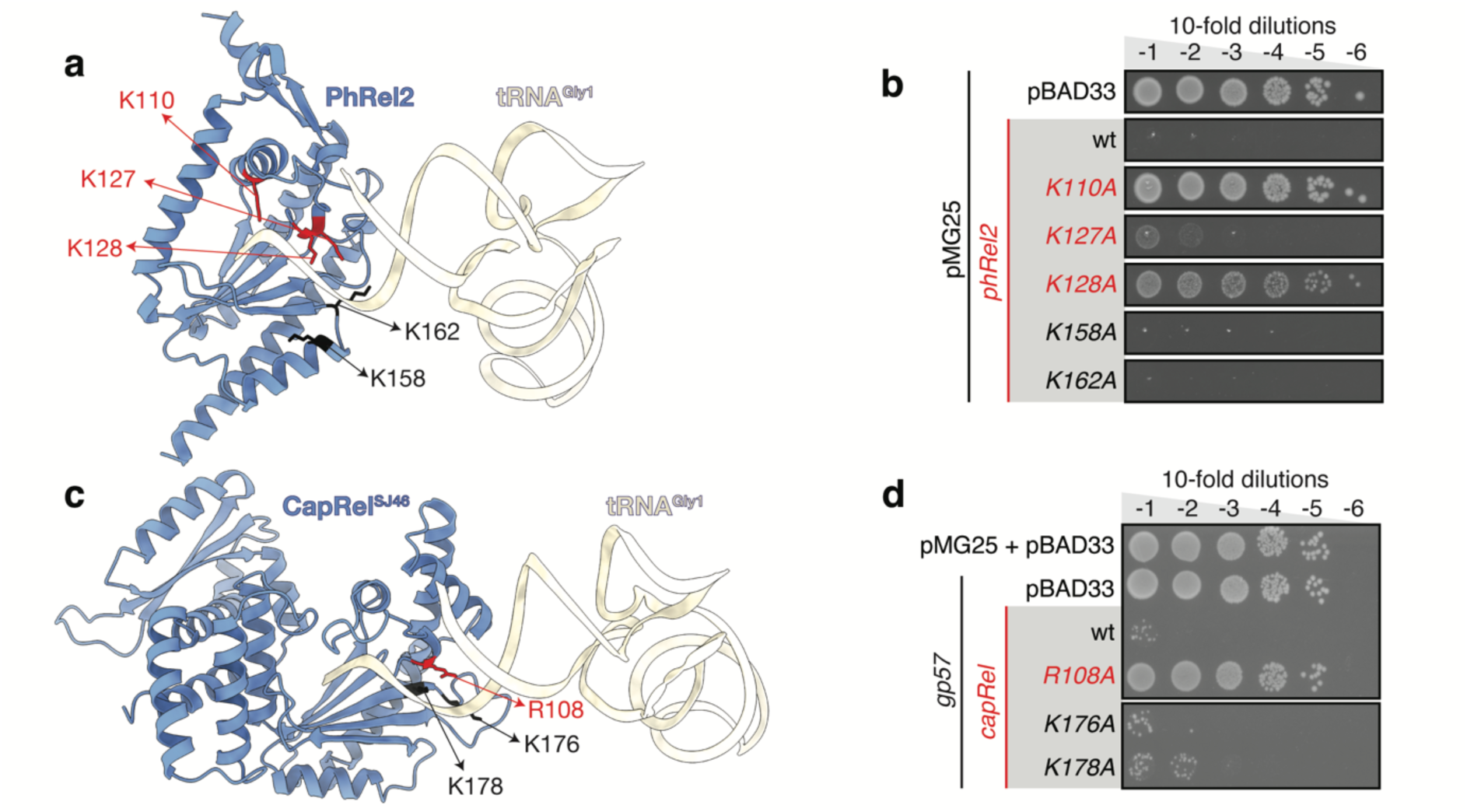
AF3-generated structures of tRNA^Gly^-bound *B. subtilis* la1a PhRel2 and CapRel^SJ46^ and their mutational probing. **(a,b)** AF3-predicted structure of *B. subtilis* la1a PhRel2 in complex with *E. coli* tRNA^Gly^ (a) and its mutational validation in toxicity assays (b). **(c,d)** Predicted structure of CapRel^SJ46^ in complex with *E. coli* tRNA^Gly^ (c) and mutational validation in toxicity assays (d). (b,d) Ten-fold dilutions of overnight cultures of *E. coli* strains transformed with pBAD33 vector or pBAD33 derivatives expressing either wild-type or mutant toxSAS variants and either pMG25 vector or pMG25 derivatives expressing Gp57 were spotted on LB plates and scored after a 16-hour-long incubation at 37°C. The experiments were performed two times, representative plates are shown.

**Supplementary Figure 7.**
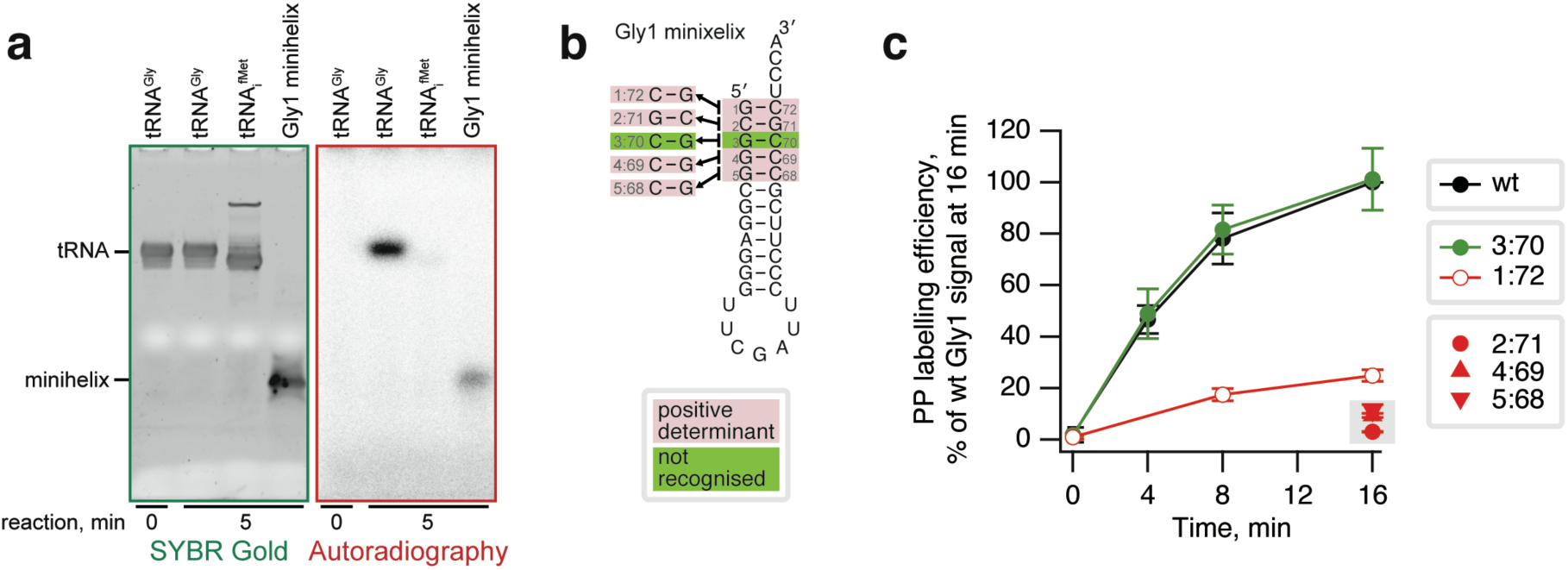
Validation of the tRNA^Gly1^-mimicking RNA minihelix as an experimental model and kinetic analysis of Gly1 minihelix (wild type and mutant variants) modification by FaRel2. (**a**) Pyrophosphorylation of native *E. coli* tRNAs as well as synthetic tRNA^Gly1^-mimicking RNA minihelix by FaRel2. The reaction mixture containing 5 µM RNA substrates, 100 µM ^32^P-labelled ATP and 5 nM FaRel2-FLAG_3_ was incubated at 37°C for 5 minutes and then quenched with RNA dye. The samples were resolved on 12% urea-PAGE and visualized by SYBR Gold staining as well as by autoradiography. The experiment was performed twice, a representative gel is shown. (**b**) Predicted secondary structure of tRNA^Gly1^-mimicking RNA minihelix as well as mutations used in kinetic experiments. (**c**) Time course of FaRel2-mediated pyrophosphorylation of synthetic Gly1 RNA minihelix substrates. 5 µM substrates were modified at 37°C for 4, 8 or 16 minutes by 10 nM FaRel2-FLAG_3_ in the presence of 100 µM ^32^P-labelled ATP, and then the reactions were quenched, RNAs resolved on 12% urea-PAGE and visualized using SYBR Gold staining and autoradiography. The autoradiography signal was normalized to SYBR Gold signal, and expressed as a faction of the signal intensity for wild-type Gly1 substrate. Signal intensity for wild-type Gly1 at 16 minutes was set to 100 %. The experiment was performed in triplicates and the quantified data is shown as average ± standard deviation.

## MATERIALS AND METHODS

### Strain and plasmid construction

*E. coli* strains, plasmids, oligonucleotides and synthetic DNA used in this study are listed in **Supplementary Table 3**. PCR-amplified DNA fragments were assembled by Gibson assembly kit (NEBuilder, NEB) as per manufacturer’s protocol and introduced into DH5α *E. coli* strain. The assembled plasmids were amplified in the DH5α cells grown in LB (Lennox) liquid media, purified using QIAprep Spin Miniprep Kit (QIAGEN) and sequenced.

### Protein expression and purification

C-terminally FLAG_3_-tagged *Coprobacillus* sp. D7 FaRel2 (FaRel2-FLAG_3_) was cloned into a pBAD33 derivative VHp678 under the control of arabinose-inducible P_BAD_ promotor. The toxin was overexpressed in *E. coli* BL21 DE3 cells co-transformed with the VHp701 plasmid encoding non-tagged Small Alarmone Hydrolase (SAH) ATfaRel antitoxin under the control of T7 promoter. Fresh transformants were used to inoculate an 800 mL culture (LB supplemented with 50 μg/mL kanamycin and 20 μg/mL chloramphenicol) to a final OD_600_ of 0.04. Bacteria were grown at 37°C until an OD_600_ of 0.3 when the antitoxin was pre-induced with 0.1 mM IPTG (final concentration) for one hour, after which the toxin was induced with 0.2% arabinose (final concentration) for an additional hour. The cells were collected by centrifugation (8,000 rpm, 10 minutes at 4°C, JLA-10.500 rotor (Beckman Coulter)), dissolved in 4 mL of cell suspension buffer (20 mM HEPES:KOH pH 7.5, 95 mM KCl, 5 mM NH_4_Cl, 0.5 mM CaCl_2_, 8 mM putrescine, 1 mM spermidine, 5 mM Mg(OAc)_2_, 1 mM DTT and cOmplete protease inhibitor (Mini, EDTA-free from Roche)). The cell suspension was divided to 1 mL aliquots, and 200 μl of pre-chilled zirconium beads (0.1 mm) were added to each of them. Cellular lysates were prepared by a FastPrep homogeniser (MP Biomedicals) (four 20 seconds pulses at speed 4.5 mp per second with chilling on ice for 2 minutes between the cycles), and clarified by centrifugation at 21,000 g for 20 minutes at 4°C. The supernatant was carefully collected, avoiding the lipid layer and the cellular pellet.

30 mg of total protein as determined by Bradford assay of each sample was combined with 100 μL of ANTI-FLAG M2 Affinity Gel (Sigma-Aldrich) and mixed by rotation for 2 hours at 4°C. The mixture was loaded on a Micro Bio-Spin Chromatography Column (Bio-Rad) and flow-through was collected. The gel in the column was washed five times with 1 mL of cell suspension buffer supplemented with 10% glycerol, and the fraction of the final wash was collected. Next, the gel was mixed for 20 min at 4°C on Multi Purpose Tube Rotator (Fisherbrand) together with 300 μL of elution buffer (cell suspension buffer additionally supplemented with 10% glycerol as well as 0.1 mg/mL Poly FLAG Peptide lyophilised powder (Biotool)). The protein was eluted by briefly spinning down the column in a table-top centrifuge and collected in Eppendorf tube. After the elution step the gel beads were resuspended with 1x sample buffer (50 mM Tris:HCl pH 6.8, 2% SDS, 0.01% bromophenol blue, 10% glycerol, 10 mM DTT and 2% beta-mercaptoethanol).

0.5 μL of cell lysate, 0.5 μL of flowthrough, 8 μL of wash, 8 μL of elution fraction as well as 10 μL of gel suspension were resolved on 15% SDS-PAGE gel. The SDS-PAGE gel was treated with fixation solution (50% methanol and 2% phosphoric acid) for 5 min at room temperature, washed three times with water for 15 minutes at room temperature and stained with “blue silver” solution^42^ (0.12% Brilliant Blue G250 (Sigma-Aldrich, 27815), 10% ammonium sulphate, 10% phosphoric acid, and 20% methanol) overnight at room temperature. After the final 3-hour wash with water for at room temperature, the gel was imaged on Amersham ImageQuant 800 imaging system (Cytiva). The concentration of FaRel2-FLAG_3_ was estimated from SDS-PAGE gels by ImageJ^43^ using pure ATfaRel2 as a standard.

### Toxicity assays

The assays were performed on LB medium (Lennox) plates (VWR). The *E. coli* BW25113 strain was transformed with pBAD33 derivatives expressing either wild-type *faRel2* or *faRel2* variants or an empty pBAD33 plasmid used as a vector control. The nucleotide sequence of the *faRel2* ORF was codon optimised for expression in *E. coli*. The cells were grown in liquid LB medium (BD) supplemented with 20 µg/mL chloramphenicol (AppliChem) as well as 0.2% glucose (repression conditions). Serial ten-fold dilutions were spotted (5 µL per spot) on solid LB plates containing chloramphenicol was well as either 0.2% glucose (repressive conditions), or 0.2% arabinose (induction conditions). Plates were scored after an overnight incubation at 37°C.

### Experimental phage infections

To assess the activity of FaRel2:ATfaRel2 TA system in phage defence, we performed efficiency of plating assays essentially as described previously^44^. The experiments were performed using the BASEL coliphage collection^28^ as well as a set of common laboratory phages. Briefly, *E. coli* BW25113 co-transformed with either VHp277 (pBAD33-*faRel2*) and VHp1199 (pMG25-*aTfaRel2*) or the empty pBAD33 and pMG25 vectors were grown overnight in LB medium supplemented with 20 µg/mL chloramphenicol and 100 µg/mL ampicillin. Importantly, the leaky *aTfaRel2* expression from VHp1199 is sufficient to neutralize the toxicity induced by *faRel2* expression^26^. Bacterial lawns were prepared by mixing 0.75 OD_600_ units of cells with 10 ml of top agar (LB with 0.2% arabinose, 0.5% agar, 20 mM MgSO_4_, and 5 mM CaCl_2_) and overlaying this mixture on square LB-agar plates (1.5% agar) containing 0.2% arabinose. Phage stocks were 10-fold serially diluted in SM buffer (100 mM NaCl, 10 mM MgSO_4_, and 50 mM Tris-HCl pH 7.5) and 2.5 μL of each of eight dilutions spotted on the solidified top agar plates. The formation of plaques was monitored after 6 and 24h of incubation at 37°C.

### anti-FLAG_3_ immunoprecipitation of FLAG_3_-tagged FaRel2

C-terminally FLAG_3_-tagged *Coprobacillus* sp. D7 FaRel2 (FaRel2-FLAG_3_) was expressed from a pBAD33-derived plasmid VHp678 under the control of P_BAD_ promotor in *E. coli* BW25113 that was co-transformed with a pMG25 derivative encoding non-tagged SAH PaSpo^SSU5^ from *Salmonella* phage SSU5 or ATfaRel2 antitoxin under the control of P_A1/O4/O3_ promoter. Fresh transformants were used to inoculate a 220 mL culture to a final OD_600_ of 0.05 in LB medium supplemented with 100 μg/mL ampicillin and 20 μg/mL chloramphenicol. Expression of the ATfaRel2 antitoxin was pre-induced with 150 µM IPTG. Expression of PaSpo^SSU5^ was not induced by IPTG as leaky expression from the P_A1/O4/O3_ promoter was sufficient to neutralise the FaRel2 toxicity. The cultures were grown at 37°C until an OD_600_ of 0.3 and then the expression of FaRel2 was induced with 0.2% arabinose (final concentration). After three hours at 37°C, the culture was divided into two: 200 mL for pulldown and 20 mL for tBulk preparation, see the corresponding section of *Methods* section. The cells were collected by centrifugation at 4,000 rpm for 10 minutes at 4°C using S-4x universal rotor (Eppendorf). Cell pellets from the 200 mL culture were dissolved in cell suspension buffer (20 mM HEPES:KOH pH 7.5, 95 mM KCl, 5 mM NH_4_Cl, 0.5 mM CaCl_2_, 8 mM putrescine, 1 mM spermidine, 5 mM Mg(OAc)_2_, 1 mM DTT and MiniEDTA-free cOmplete protease inhibitor (Roche)) to final OD_600_ of 200. The cell suspension was divided to 1 mL aliquots, and 200 μL of pre-chilled zirconium 0.1 mm beads were added to each aliquot. Cellular lysates were prepared using FastPrep homogeniser (MP Biomedicals) via four 20-second pulses at speed 4.5 mp per second with chilling on ice for 2 minutes between the cycles. The lysates clarified by centrifugation at 21,000 g for 20 minutes at 4°C and the supernatant was carefully collected avoiding the lipid layer and the cellular pellet.

Protein concentration in the supernatant was determined by Bradford assay, 5 mg of total protein per sample was combined with 100 μL of ANTI-FLAG M2 Affinity Gel (Sigma-Aldrich) and mixed on the end-to-end rotator for 2 hours at 4°C. The mixture was loaded on a Micro Bio-Spin Chromatography Column (Bio-Rad), the flow-through was collected and kept for further analysis. The column was washed five times with 1 mL of cell suspension buffer, and a fraction at final wash was collected. Using an end-to-end rotator, the gel was mixed on the column for 20 min at 4°C with 300 μL of cell suspension buffer supplemented with 0.1 mg/mL Poly FLAG Peptide (Biotool). The sample was eluted from the column by centrifugation and was collected in Eppendorf tube. In total 900 µL of eluate was collected, and three technical replicates were performed for each biological replicate. After the elution step, the gel beads were suspended with 1x sample buffer (50 mM Tris:HCl pH 6.8, 2% SDS, 0.01% bromophenol blue, 10% glycerol, 10 mM DTT and 2% beta-mercaptoethanol) and kept for further analysis. 0.5 μL of cell lysate, 0.5 μL of flowthrough, 8 μL of wash, 8 μL of elution fractions and 10 μL of gel suspension were resolved on 12% SDS-PAGE gel. The SDS-PAGE gel was imaged. The FaRel2-FLAG_3_ concentration was estimated as described in the *Protein expression and purification* section.

The eluted sample (900 µL) was mixed with an equal volume of acidic phenol pH 4.3 and centrifugated (14,000 rpm for 20 minutes at 4°C in 5418 R Centrifuge equipped with FA-45-18-11 rotor (Eppendorf)). The aqueous phase was collected, transferred to a new Eppendorf tube, mixed with equal volume of chloroform and the two phases were separated by centrifugation (14,000 rpm for 1 minute at 4°C). The aqueous phase was again transferred to a new Eppendorf tube, mixed with 1/100 volume of 2 mg/mL glycogen, 3/50 volume of 5 M NaCl_2_ and 2.5 volume of 96% EtOH and kept at-20°C for overnight. Next, the tRNA sample was precipitated by centrifugation (14,000 rpm for 30 minutes at 4°C) and the pellet was washed with 200 µL of 70% EtOH. After air-drying at room temperature for 5 minutes, the pellet was dissolved in 10 µL of nuclease-free water.

Do dephosphorylate the purified tRNA preparations, 9.5 µL of tRNA was combined with 2 U/µL of T4 PNK (NEB, M0201S) was well as N-terminally His_6_-TEV-tagged ATfaRel SAH (final concentration 1 µM) in a 20-µL reaction mixture (1x Polymix buffer with 5 mM Mg^2+^ final concentration additionally supplemented with 1 mM MnCl_2_ and 1 mM DTT). After a 10-minute incubation 37°C, the reaction was stopped by addition of 40 µL of RNA loading dye (98% formamide, 10 mM EDTA, 0.3% BPB and 0.3% Xylene cyanol). A 60 µL sample was resolved on 8 M urea-PAGE/TBE (8% acrylamide/bis-acrylamide = 19:1); 20 µL of the sample was loaded per lane. After staining the tRNA with SYBR Gold, the gel pieces containing tRNA were cut out and crushed in 1.5 mL Eppendorf tube. The crushed gel was combined with 1.2 mL of elution buffer (0.3 M NaOAc pH5.2, 0.1% SDS, and 1 mM EDTA), and after shaking at 1500 rpm for 2 hours at 37°C, the supernatant was separated by passing the mixture through a 0.22 µm filter. The elution step was repeated one more time with fresh elution buffer and collected tRNA elution samples were pooled. The pooled samples were mixed with 1/100 volume of 2 mg/mL glycogen and 2.5 volume of 96% EtOH, and then kept at-20°C overnight. tRNA solution was aliquoted in 1.5 mL tubes, and tRNAs were pelleted by centrifugation (14,000 rpm for 30 minutes at 4°C). The pellets were washed with 200 µL 70% EtOH and after air-drying pellet at room temperature for 5 minutes, the resultant tRNA preparations were kept at-80 °C until use. To assess the tRNA yield and quality, the pellet was dissolved with nuclease free water, the concentration was measured spectrophotometrically assuming 1 A_260_ = 40 µg/mL.

### Preparation of E. coli tBulk

20 mL *E. coli* culture was prepared as described in the *anti-FLAG_3_ pulldown and tRNA isolation* section. Cells were collected by centrifugation (4,000 rpm, 10 minutes at 4°C, S-4x universal rotor (Eppendorf)). The cell pellets were dissolved in 400 µL of acidic cell suspension buffer (50 mM NaOAc and 10 mM Mg(OAc)_2_, pH 5.2) and mixed with 400 µL of acid phenol pH 4.3 for 5 minutes at room temperature. The mixture was frozen in liquid nitrogen and thawed in water at room temperature. After one more round of this freeze and thaw, the sample was mixed by rotation for 2 h at room temperature. Aqueous phage was separated by centrifugation (14,000 rpm for 10 minutes at room temperature in 5418 R Centrifuge equipped with FA-45-18-11 rotor (Eppendorf)), collected into new tube and mixed with 1 volume of TRI Reagent solution (Invitrogen, AM9738) and 0.1 volume of 1-bromo-3-chloropropane. The mixture was centrifugated (14,000 rpm for 10 minutes at room temperature), and aqueous phase was collected into fresh 1 mL tube. The aqueous phase was mixed with 1/20 volume of 3 M NaOAc and 1 volume of isopropanol, and then kept at-20°C for 20 minutes. Total RNA was precipitated by centrifugation (14,000 rpm for 12 minutes at 4°C), supernatant was discarded. The RNA pellet was dissolved in 300 µL nuclease-free water, mixed with 1/10 volume of 3 M NaOAc pH5.2 and 2.5 volume of 96% EtOH, and then kept at-20°C for 20 minutes. Total RNA was precipitated by centrifugation (14,000 rpm for 12 minutes at 4°C), the pellet was washed with 1 mL 70% EtOH. After air-drying pellet at room temperature for 5 minutes, the pellet was dissolved with 10 µL of nuclease free water. was washed with 200 µL 70% EtOH. After air-drying pellet at room temperature for 5 minutes, the pellet was dissolved with 7 µL of nuclease free water. Total RNA solution was mixed with 14 µL of RNA loading dye (98% formamide, 10 mM EDTA, 0.3% BPB and 0.3% Xylene cyanol), 21 µL sample (in a lane) was resolved on 8 M urea-PAGE/TBE (8% acrylamide/bis-acrylamide = 19:1). After staining tRNA with SYBR Gold, the gel pieces containing tRNA bands were cut out and mashed in 1.5 mL tube. The mashed gel was mixed with 200 µL of 10 mM Tris:HCl pH7.5 at 1500 rpm for 2 hours at 37°C, the supernatant was collected with passing through 0.22 µm filter. This elution step was repeated one more time with fresh 10 mM Tris:HCl pH7.5, collected elution containing tBulk was pooled. The pooled elution was mixed with 1/100 volume of 2 mg/mL glycogen, 3/50 volume of 5 M NaOAc and 2.5 volume of 96% EtOH, and then kept at-20°C for overnight. tBulk was pelleted by centrifugation (14,000 rpm for 30 minutes at 4°C), the pellet was washed with 200 µL 70% EtOH. After air-drying pellet at room temperature for 5 minutes, the pellet was dissolved in 10 µL nuclease free water, and the concentration was measured A_260_ value (1 A_260_ = 40 µg/mL). Ten micrograms tBulk was dephosphorylated and re-purified as described in the *anti-FLAG_3_ pulldown and tRNA isolation* section.

### Expression and purification of native glycine-specific tRNA^Gly^

The native tRNA was expressed and purified essentially as described previously, ref.^45^ In brief, *E. coli* tRNA^Gly1^ (CCC anticodon) genomic sequence was assembled from five oligonucleotides, supplemented with flanking *Eco*RI and *Pst*I restriction sites, see **Supplementary Table 3**.

The assembled tRNA genomic sequence was then inserted into the *Eco*RI/*Pst*I-digested pBSTNAV vector (AddGene, USA). The final construct contained the tRNA under the control of the constitutive lpp promoter. The cells were grown overnight (16h) in LB media supplemented with 100 µg/ml ampicillin, and the biomass was collected by centrifugation. It was resuspended in 1 mM Tris-HCl pH 7.4, 10 mM Mg(CH_3_COO)_2_ and lysed with 0.5 volumes of acidic phenol:chloroform mix 5:1 pH 4.5. The aqueous phase was precipitated and resuspended in 1M NaCl. The solution, containing soluble RNAs was precipitated again. To deacylate bulk tRNA, the pellet was resuspended in 200 mM Tris-HCl pH 9.0 and incubated for 2h at 37°C. The deacylated tRNA was ethanol precipitated and dissolved in monoQ buffer A (40 mM sodium phosphate buffer, pH 7.0) and separated on the 8-ml MonoQ column (10/100, GE Healthcare), using two-step linear gradient of buffer B (A with 1M NaCl): 20 ml 0-50%B followed by 280 ml 50-100%. The tRNA^Gly1^-containing fractions were identified by analytical aminoacylation with [^14^C]-glycine using *Thermus thermophilus* GlyRS. Pooled fractions were precipitated and dissolved in C5 buffer A (20 mM NH_4_CH_3_COO pH 5.5, 400 mM NaCl, 10 mM MgCl_2_, 1 mM EDTA, 1% methanol). Further reverse-phase chromatography on the C5 column (C5-5, 250 × 10 mM, Discovery BIO Wide Pore, Supelco) using 300-ml 0-60% linear gradient of C5 buffer B (Buffer A supplemented with 40% methanol) produced pure tRNA preparation with over 95% glycine-charging activity.

### mim-tRNAseq tRNA sequencing and data analysis

mim-tRNAseq was performed as described previously^31^ with minor modifications. Seven picomols of tBulk and FaRel2-bound tRNA prepared from *E. coli* cells co-expressing FaRel2-FLAG_3_ with either ATfaRel2 or PaSpo^SSU5^ were deacylated under 100 μL of 100 mM CHES-NaOH pH 9.0 for 1 hour at 37°C and purified by Oligo Clean & Concentrator (Zymo Research). Before adapter ligation, DNA adapter was adenylated in a 30 μL reaction mixture consisting of 6 μM DNA adapter, 5 μM Mth RNA ligase (NEB), 1 x adenylation buffer (NEB), and 1 mM ATP, and purified by Oligo Clean & Concentrator. The deacylated tRNA was ligated with the DNA adapter to the 3’ end in a 20 μL reaction mixture consisting of 28 pmol adenylated DNA adapter, 200 U T4 RNA ligase 2 truncated KQ (NEB), 1x T4 RNA ligase buffer (NEB), 25% PEG8000, and 10 U SUPERase•In (Thermo Fischer Scientific) at 22°C overnight. After ligation, the mixture was purified by Oligo Clean & Concentrator and resolved on a urea-PAGE in 1 x TBE (7 M urea, 10% PAGE). The gel was stained with SYBR Gold (Thermo Fischer Scientific) and gel-image was acquired by a FAS-Digi PRO (NIPPON Genetics). Visualized bands corresponding to adapter-ligated tRNAs were excised from the gel and eluted for 3 h at 37°C with continuous mixing in 400 μL of elution buffer consisting of 400 mM NaOAc pH 5.2, 0.1% SDS, and 1 mM EDTA-NaOH pH8.0. The elute was filtered by Ultrafree-MC (Merck) to remove the pieces of mashed gel and then subjected to ethanol precipitation with 20 μg/mL glycogen. For reverse transcription, adapter-ligated tRNA was mixed with 2.5 pmol RT primer in a 11 μL water, denatured at 82°C for 2 minutes, and annealed at 25°C for 5 minutes, and then applied to reverse transcription at 42°C overnight in a 20 μL reaction mixture consisting of 1x RT buffer (50 mM Tris-HCl pH 8.3, 75 mM KCl, and 3 mM MgCl_2_), 5 mM DTT, 5 mM dNTP, 20 U SUPERase In, and 1 μL TGIRT-III (InGex). Following reverse transcription, tRNA template was hydrolysed by alkaline treatment with 100 mM NaOH at 95°C for 2 minutes and synthesized cDNA was purified by Oligo Clean & Concentrator, followed by gel purification as described above. Obtained cDNA was circularized at 60°C overnight in a 20 μL reaction mixture consisting of 10 U with CircLigase ssDNA ligase (Lucigen), 1x reaction buffer (Lucigen), 2.5 mM MnCl_2_, 1 M betaine and 50 μM ATP, and incubated at 80°C for 10 minutes. One micro litter of circularized cDNA was subjected to PCR in a 20 μL reaction mixture composed of 20 U/mL Phusion polymerase (NEB), 1x Phusion GC buffer (NEB), 0.2 mM dNTPs, 3% DMSO, 0.5 μM forward primer, and 0.5 μM reverse primer. The reaction was performed with initial denaturation at 98°C for 30 sec, followed by 8 or 10 cycles of 98 °C for 10 sec, 60°C for 20 sec, 72°C for 5 sec. PCR product was purified by AMPure XP (Beckman Coulter) and further purified by gel extraction with native-PAGE in 1x TBE (6% PAGE). Obtained cDNA libraries were purified by AMPure XP and quantified by Agilent 2100 Bioanalyzer system (Agilent). 500 pmol of each cDNA library was mixed in a 30 μL water and sequenced on an Illumina HiSeq X Ten platform (150 bp, pair end).

Raw Illumina sequencing reads were trimmed to remove both adapter and Read 1 or 2 sequences using cutadapt 4.4 (default version). Only Read 1 sequence data were used for subsequent processing. Following trimming, 17 nt of random UMI sequence (N14D3) attached to 5’ side of tRNA sequence was further removed from each read and written into the read name as a UMI tag by fastp v0.23.2. The reads with shorter than 15 nt and quality score at 5’ and 3’ ends below 20 were discarded by trimmomatic-0.39, and the filtered reads were then aligned to the sequences of *E. coli* non-coding RNAs including all tRNAs (obtained from NCBI database (*E. coli* BW25113 complete genome, GenBank CP009273.1) using bowtie 2-2.5.1-linux-x86_64 ^46^ with very sensitive local mode and-L 10. PCR duplicates were deduplicated based on the UMI tag by UMICollapse. The mapped read numbers on each tRNA gene were counted by samtools-1.14. The *p*-value was calculated by two-sided Student’s *t*-test, n = 3. Statistics of the individual tRNA coverage (read counts) for mim-tRNAseq experiments are provided as a **Supplementary Table 1**.

### tRNA Northern blotting

We used *E. coli* BW25113 strain transformed either with pBAD33 derivative expressing wild-type *faRel2* or empty pBAD33 as vector control. The cells were grown in liquid LB medium (BD) supplemented with 20 µg/mL chloramphenicol (AppliChem) as well as 0.2% glucose (to repress the FaRel2 expression). Fresh transformants were used to inoculate a 40-mL culture (LB supplemented with 20 μg/mL chloramphenicol) to a final OD_600_ of 0.05. Bacteria were grown at 37°C until an OD_600_ of 0.3, and then FaRel2 expression was induced for 30 min with 0.2% arabinose (final concentration). The cells from a 25-mL culture were collected by centrifugation (4,000 rpm, 10 min, at 4°C in S-4x universal rotor (Eppendorf)) and resuspended in 0.5 mL of 3 M NaOAc pH4.5 supplemented with 10 mM EDTA. The cells mixed with 0.5 mL of acid phenol and 20 µL BCP and keept on ice for 15 min. After centrifugation (14,000 rpm for 20 min at 4°C in 5418 R Centrifuge equipped with FA-45-18-11 rotor (Eppendorf)), the aqueous phage was removed and mixed with an equal volume of isopropanol and kept at –20°C for an hour. Total RNA was precipitated by centrifugation (14,000 rpm for 20 min at 4°C) and the RNA pellet was washed with 1 mL 70% ethanol. The RNA pellet was dissolved in 20 µL of 10 mM NaOAc pH4.5 supplemented with 1 mM EDTA, total RNA concentration was determined spectrophotometrically (1 A_260_ = 40 µg/mL). The RNA sample was mixed with ≥5 volumes of acid urea-PAGE sample buffer (98% formamide, 10 mM EDTA, 0.3% BPB and 0.3% Xylene cyanol for nucleic acid staining) and 6 µg RNA was resolved on 8 M urea-PAGE/100 mM NaOAc pH5.2 (6.5% acrylamide:bis-acrylamide in 19:1 ratio). RNA was transferred to Zeta-Probe^®^ Blotting Membranes (Bio-Rad) using Trans-Blot^®^ TurboTM Transfer System (Bio-Rad), UV cross-linked on menbrane, and the membrane was pre-hybridized at 42°C for ≥3 h in pre-heated Church buffer (0.25 mM Na_2_HPO_4_, 0.17% orthophosphoric acid, 1 mM EDTA, 0.01% bovine serum albumin and 0.07% SDS). After discarding the used Church buffer, 13 nM ^32^P-labeled probe was hybridized in fresh Church buffer for ≥16 h at 42°C. The membrane was 3 times-washed with 0.1% SDS/6x SSC at 42°C for 5 min, exposed on an imaging plate and the plate was imaged by a FLA-3000 (Fujifilm).

### tRNA pyrophosphorylation assays

With the exception of the experiment shown on **Figure 3b**, FaRel2-FLAG_3_-mediated tRNA modification was assayed in two enzymatic regimes: i) low enzyme concentration (5-10 nM) combined with high substrate concentration (5 µM) and ii) high enzyme concentration (50-100 nM) combined with low substrate concentration (0.4 µM).

FaRel2-FLAG_3_ was reacted with either i) co-immunoprecipitated tRNA, ii) tBulk purified from *E. coli* as described in the section *Preparation of* E. coli *tBulk*, see above, iii) commercial tBulk from Roche (tBulk_Roche_) or iv) *E. coli* tRNA_i_^fMet^, tRNA^Phe^, tRNA^Val^ (all from Chemical Block) as well as tRNA^Gly^ (purified as described in the section *Expression and purification of native glycine-specific tRNA^Gly^*, see above). Conditions for experiments presented on different figures are as follows:

**Figure 3b:** toxSAS 50 nM, tRNA substrate 5 µM, reaction time 3-10 min

**Figure 3c:** toxSAS 50 nM, tRNA substrate 0.4 µM, reaction time 10 min

**Figure 5c:** toxSAS 5 nM, tRNA substrate 5 µM, reaction time 5 min

**Figure 5d:** toxSAS 10 nM, tRNA substrate 5 µM, reaction time 4-16 min

**Supplementary Figure 4**: toxSAS 100 nM, tRNA substrate 0.4 µM, reaction time 10 min

**Supplementary Figure 7a**: toxSAS 5 nM, tRNA substrate 5 µM, reaction time 5 min

**Supplementary Figure 7c**: toxSAS 10 nM, tRNA substrate 5 µM, reaction time 4-16 min.

The reactions were started by the addition of 500 µM γ^32^P-ATP and carried out at 37°C for either 0.25, 3, 5 or 10 minutes. To visualise pyrophosphorylated tRNA the reaction sample was mixed in 2 volumes of RNA dye (98% formamide, 10 mM EDTA, 0.3% bromophenol blue and 0.3% xylene cyanol), tRNA denatured at 37°C for 10 min and resolved on urea-PAGE in 1 x TBE (8 M urea, 8% PAGE). The gel was stained with SYBR Gold (Life technologies, S11494) and imaged using Amersham™ ImageQuant^TM^ 800. Next, the gel was exposed to an imaging plate overnight and the imaging plate was imaged by a FLA-3000 (Fujifilm). In the case of kinetically-resolved experiments, the signal intensities from the SYBR Gold staining and the phosphorimaging were quantified using ImageJ^43^ software. The data was processed as follows: i) the relative loading amounts of individual minihelix species were calculated as ratio of averaged signals from SYBR gold staining of individual loading replicates, ii) individual phosphorimaging signal intensities were normalised to the relative loading amount of the minihelix species, iii) efficiency of the pyrophosphoate labelling was calculated as percentile to the normalised intensity of wild-type at 16 minutes (set to 100%), and iv) the average and the standard deviation were calculated using the data from three independent experiments.

### Cell-free translation assays

Experiments with PURExpress In Vitro Protein Synthesis Kit (NEB, E6800) were performed as per the manufacturer’s instructions using 10 ng/µL of DHFR-Strep reporter plasmid with the addition of 0.8 U/µL RNase Inhibitor Murine (NEB, M0314S). Purified FaRel2-FLAG_3_ (at 50 nM as final concentration) or PhRel2-FLAG_3_ (at 100 nM as final concentration) was used, and as a mock control, toxSASes was substituted for equal volume of HEPES:Polymix buffer. The total reaction volume was 6 µL per reaction for most of the experiments. After incubation at 37°C for the indicated time, the reaction mixture was mixed with 9-fold volume of 2x sample buffer (100 mM Tris:HCl pH 6.8, 4% SDS, 0.02% bromophenol blue, 20% glycerol, 20 mM DTT and 4% β-mercaptoethanol), and 5 µL of the mixture was resolved on 18% SDS-PAGE gel. In the case of CapRel^SJ46^, purified CapRel^SJ46^ protein was used at a final concentration of 250 nM, with *gp57*, an essential activator for CapRel^SJ46^, as template plasmid at 10 ng/µL. As a mock control CapRel^SJ46^ was substituted for equal volume of HEPES:Polymix buffer. After a 10-minute incubation at 37°C, a 1.34 µL aliquot of the reaction mixture was taken and quenched by addition of 13.66 µL of 2x sample buffer, and DHFR-Strep reporter plasmid solution (193 ng/µL) was added to the remaining reaction mixture at a final concentration of 20 ng/µL. After further incubation at 37°C for 1 hour, the reaction mixture was mixed with 9-fold volume of 2x sample buffer and 5 µL of the mixture was resolved on 18% SDS-PAGE gel. The SDS-PAGE gel was fixed by incubating for 5 min at room temperature in 50% methanol solution supplemented with 2% phosphoric acid, then stained and detected as mentioned in protein expression and purification.

### Sucrose gradient fractionation and Western blotting

C-terminally FLAG_3_-tagged *Coprobacillus* sp. D7 FaRel2 variant (FaRel2-FLAG_3_) was expressed from arabinose-inducible P*_BAD_* promotor on pBAD33-derivative VHp1093 in *E. coli* BW25113 cells co-transformed with the VHp847 plasmid encoding Salmonella phage SSU5 SAH PaSpo under the control of IPTG-inducible P*_A1/04/03_* promoter. Fresh transformants were used to inoculate a 100-mL culture (LB supplemented with 100 μg/mL ampicillin and 20 μg/mL chloramphenicol) to a final OD_600_ of 0.05. Bacteria were grown at 37°C until an OD_600_ of 0.3 without IPTG, and then FaRel2-FLAG_3_ K28A R29A was induced with 0.2% arabinose (final concentration) for an hour. The cells from 50 mL culture were collected by centrifugation (4,000 rpm, 10 min, at 4°C in S-4x universal rotor (Eppendorf)), frozen in liquid nitrogen and stored at –80°C until use. The cells were melted on ice and dissolved in 0.5 mL of HEPES:Polymix buffer (ref.^47^, 5 mM Mg^2+^ final concentration) supplemented with 1 mM PMSF, lysed using FastPrep homogenizer (MP Biomedicals) (four 20 second pulses at 4.0 m/s with chilling on ice for 2 min between the cycles), and clarified by centrifugation (14,000 rpm for 20 min in 5418 R Centrifuge equipped with FA-45-18-11 rotor (Eppendorf)). 10 A_260_ units of the lysate was loaded onto 10-35% sucrose gradient in HEPES:Polymix buffer pH 7.5 (5 mM Mg^2+^ final concentration) and resolved by ultracentrifugation at 36,000 rpm for 3 h at 4°C with the fastest braking in Optima XPN-80 Ultracentrifuge equipped with SW-41Ti rotor (Beckman Coulter). Gradients were fractionated (0.5 mL/fraction) using Biocomp Gradient Station (BioComp Instruments) with A_260_ as a readout. For Western blotting, 0.5 mL fractions were supplemented with 1.5 mL of 96% ethanol and precipitated overnight at –20°C. After centrifugation at 14,000 rpm for 30 min at 4°C, the supernatants were discarded. and the samples were dried. The pellets were resuspended in 40 µL of 1x SDS loading buffer (50 mM Tris-HCl pH 6.8, 2% SDS (w/v) 0.01% Bromophenol blue, 10% glycerol (w/v) and 2% β-mercaptoethanol), resolved on the 10% SDS PAGE and transferred to nitrocellulose membrane (Trans-Blot Turbo Midi Nitrocellulose Transfer Pack, Bio-Rad, 0.2 µm pore size) with the use of a Trans-Blot Turbo Transfer Starter System (Bio-Rad). Membrane blocking was done for 1 h in PBS-T (1xPBS 0.05% Tween-20) with 5% w/v non-fat dry milk at room temperature. FaRel2 was detected using anti-FLAG M2 antibody (Sigma-Aldrich, F1804; 1:10,000 dilution) primary, combined with Goat anti-Mouse IgG-HRP (Agrisera, AS111772; 1:5,000 dilution) as well as PBS-T. ECL detection was performed using WesternBright^TM^ Quantum (K-12042-D10, Advansta) Western blotting substrate and Amersham™ ImageQuant^TM^ 800 imaging system (Cytiva).

### Structural modeling

The structures of tRNA^Gly1^-bound FaRel2, PhRel2 and CapRel^SJ^^46^ in the ATP-liganded state were predicted using AlphaFold 3 (AF3), ref.^30^, via AlphaFold Server (https://alphafoldserver.com/).

## Data availability

The tRNA sequencing data generated in this study have been deposited on National Center for Biotechnology Information (NCBI’s) sequence read archive (SRA) database under accession code PRJNA1090630.

